# Repertoire-behavior mapping reveals signal functions in cooperatively breeding crows

**DOI:** 10.64898/2026.04.02.715916

**Authors:** Maddie Cusimano, Benjamin Hoffman, Carlos Guzón-García, Angela Mcenerey, Margaret Bezrutczyk, Jen-Yu Liu, Logan S. James, Sara C. Keen, Louis Mahon, Olivier Pietquin, Milad Alizadeh, Damián E. Blasi, Matthieu Geist, Marius Miron, David Robinson, Marta Vila, Eva Trapote, Christian Rutz, Emmanuel Chemla, Daniela Canestrari, Vittorio Baglione

## Abstract

Communication structures society, and is likewise shaped by relationships and shared tasks; yet, for most socially complex species, we know little of their full vocal repertoire and its functions. We investigated how communication structures the who, what, and when of social interactions in cooperative carrion crows – group-living birds who rely on coordinated behaviors, as in chick care. Leveraging machine learning to integrate large-scale data from crow-borne audio-loggers and nest cameras, we charted the vocal repertoire across 24 cooperative groups and mapped all discovered call types to behaviors and social context. We found that crows used a rich repertoire across three domains of joint behavior – flocking, chick care, and territorial display. Relatively quiet call types were abundant and included close-range calls that may coordinate chick care by announcing nest visits. Our study demonstrates how combining continuous-capture data and machine learning can reveal a holistic understanding of how vocalizations function across contexts.

## 1 Introduction

Group-living animals communicate for many reasons, including to coordinate movement (meerkats, Demartsev, Averly, et al., 2024), alert others to threats (scrub jays, McGowan and Woolfenden, 1989), share the location of food sources (honey bees, Gould, 1974), and increase social cohesion (naked mole rats, Barker et al., 2021). Considering particular behaviors or signals in isolation can hide complexity in communication systems by obscuring patterns that emerge across contexts (Price et al., 2015; Elie and Theunissen, 2018). Continuously-recorded observations of everyday animal behavior across contexts can now be processed at scale by machine learning (Tuia et al., 2022; Stowell, 2022; Hoffman et al., 2024), raising the possibility of systematically mapping components of complex signal repertoires to diverse behaviors and environmental conditions (Rutz, Bronstein, et al., 2023). Here we developed such an approach to investigate vocal communication in a wild bird with a facultative cooperative breeding behavior, the carrion crow *Corvus corone* (henceforth “crows”).

Unlike elsewhere in Europe, crows in northern Spain live in a complex society, where birds within a social group cooperate to build nests, raise chicks, and defend their territory (Baglione, Canestrari, Marcos, Griesser, et al., 2002). It seems likely that their well-coordinated behaviors (Trapote et al., 2024) are accompanied by an equally rich communication system (Leighton, 2017), providing a valuable opportunity to study communication in a prosocial, alloparenting animal. Yet, despite previous reports that crows not only caw, but also whistle, rattle and more (Siriwardena, 1995), and recent demonstrations that they can learn to count aloud (Liao et al., 2024), their vocal repertoire remains largely uncharacterized (Baglione, Canestrari, Cusimano, et al., 2025). More importantly, the function of carrion crow vocalizations, and the social and ecological factors shaping their usage, are still largely unknown (Wascher and Reynolds, 2025).

## 2 Results

To explore vocalization function across the crow repertoire, we collected a large-scale “dayin-the-life” dataset of vocalizations and behaviors for 51 crows across 24 social groups, while they cooperatively raised their chicks (Tables S1-S2). Video cameras at nests recorded visits by adults, who were identified by wing tags (Figure 1A), while low-impact miniature tail-mounted tags on 43 individuals recorded sounds and body movement for up to six days each (Figure 1B) (Baglione, Canestrari, Cusimano, et al., 2025; Demartsev, Gersick, et al., 2023; Stidsholt et al., 2019; Rutz and Troscianko, 2013). We used supervised machine learning to detect flights and vocalizations in audio and to attribute these vocalizations to specific individuals (Figure 1B) (Mahon et al., 2025; Denton, Wisdom, and Hershey, 2022; Hagiwara, 2023; Chen et al., 2022), and signal processing to determine the activity level (low or high) of non-flight/non-visit periods from accelerometry (Wilson, Börger, et al., 2020; Elliott, 2025) (Figure 1B). The final dataset comprised 114,676 adult crow vocalizations, 6,057 nest visits, and 36,100 flights, synchronized within a social group (Figure 1C). We could then: (a) comprehensively characterize the vocal repertoire; (b) discover interpretable associations between vocalizations and behaviors of signallers and receivers, considering daily context, group composition, and the signaller’s social category (Figure 1D, middle); and (c) interpret these results in light of the crows’ society (Figure 1D, right).

**Figure 1.**
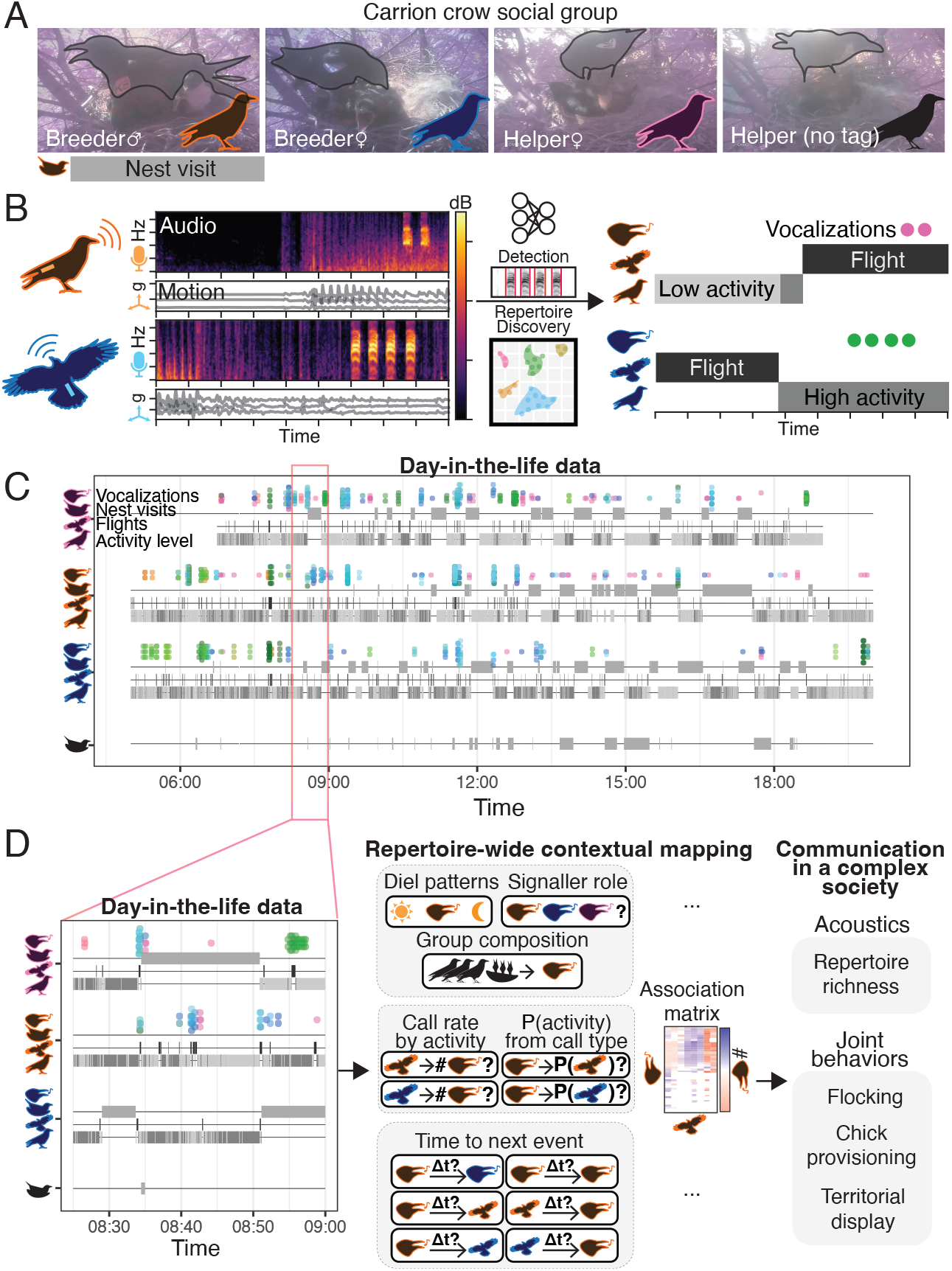
Repertoire-wide contextual mapping to characterize communication in a cooperative bird. A) Cooperative carrion crow social group. Nest camera video showed when adult crows visited the nest to care for chicks. Not all group members were tagged. B) Loggers worn by crows recorded sound and body movement. Machine learning was used to detect flights and vocalizations from audio, and guide the discovery of the vocal repertoire. Accelerometry gave the activity level (low or high) when the adult birds were neither flying nor at the nest. C) Vocalizations, nest visits, flights, and low- or high-activity periods, were synchronized into a timeline for the birds in a group. Dots: vocalizations (colors: call types, see Figure 2; stacking: vocalizations fall within a time window). Bars: behaviors. D) Four mapping analyses were applied to the timeline data (1: diel patterns, group composition, and signaller role; 2: call rate by activity; 3: P(activity) from call type; 4: time to next event). These analyses were organized into association matrices. Results were integrated to understand communication within crow society, by examining repertoire richness and its role in joint behaviors.

First, in line with the social complexity hypothesis (Leighton, 2017; Freeberg, Dunbar, and Ord, 2012), we predicted that the crows would have a rich vocal repertoire (Figure 1D, top-right). In particular, we predicted that quiet vocalizations, suggestive of close-range communication, would comprise a diversity of call types: such calls have been informally described across corvid species (Marzluff and Angell, 2013) but are difficult to measure with traditional methods (Baglione, Canestrari, Cusimano, et al., 2025). Second, we examined communication in the context of known joint activities: flocking, cooperative chick care, and cooperative territorial display (Figure 1D, bottom-right). We predicted that a set of calls would announce and follow nest visits, which could facilitate adults’ coordinated turn-taking during chick care (Trapote et al., 2024).

### 2.1 Repertoire-wide contextual mapping

We took a hybrid approach to discover the crow repertoire, using machine learning-based clustering to initially organize the large vocalization dataset and thereby make more precise manual annotation of call types tractable. Call types were generally produced by a large number of individuals, with several categories showing high effective numbers of contributors and relatively even usage across the population (Table S3). Although some call types exhibited more uneven distributions, they were nonetheless shared among multiple callers. Overall, these patterns indicate that our classification captures broadly shared vocal categories rather than reflecting primarily individual-specific vocal variants.

We then systematically interpreted each call type in terms of three levels of contextual detail, controlling for individual and group identity (Figure 1D, middle; for full statistical procedures, see Methods). At the broadest scale, we investigated which social factors (e.g., social category: breeding male, breeding female, or helper; group composition) and hours of day were associated with variation in vocalization usage (Analysis 1: Poisson GLMM to predict call rate). Narrowing in, we explored how ongoing activities (e.g., flying, staying at nest) were associated with different call types (Analysis 2: Poisson GLMM to predict call rate from activity; Analysis 3: Gaussian LMM to predict activity probability from call type). Finally, at the finest scale, we examined when events (e.g., taking off, vocalization bout) preceded, or followed, various vocalization types (Analysis 4: Gaussian LMM to predict the time between one event type and another). We performed Analyses 2–4 within a single bird (the signaller) and across two different birds in the same group (the signaller and a potential receiver) when their events were synchronized. This resulted in several association matrices between vocalizations, signaller’s behaviors, and others’ behaviors, comprising a comprehensive, yet structured, characterization of functional associations (Figures S1-S7 show the full results).

### 2.2 Acoustic complexity

As hypothesized, we found that the crows have a rich vocal repertoire. We organized the repertoire into 40 call types across three super-categories termed exceptional, caws, and grunts (Figure 2A/B; listen to examples online at https://projects.earthspecies.org/cooperative-crows/repertoire_examples.html). The exceptional super-category was manually divided into 22 diverse types (Figure 2A, left), including trills (E72), harmonic calls combined with clicks (E51), long whistles (E14), and time-varying amplitude modulated calls (E74). Exceptional calls tended to be rare, other than E4 (Figure 2B), and the only six social-category specific vocalizations were exceptional calls (Analysis 1; pairwise contrasts with *p*_BH_ < 0.05, using Benjamini-Hochberg corrections for multiple comparisons; Figure 2B), suggesting that these have specialized functions.

**Figure 2.**
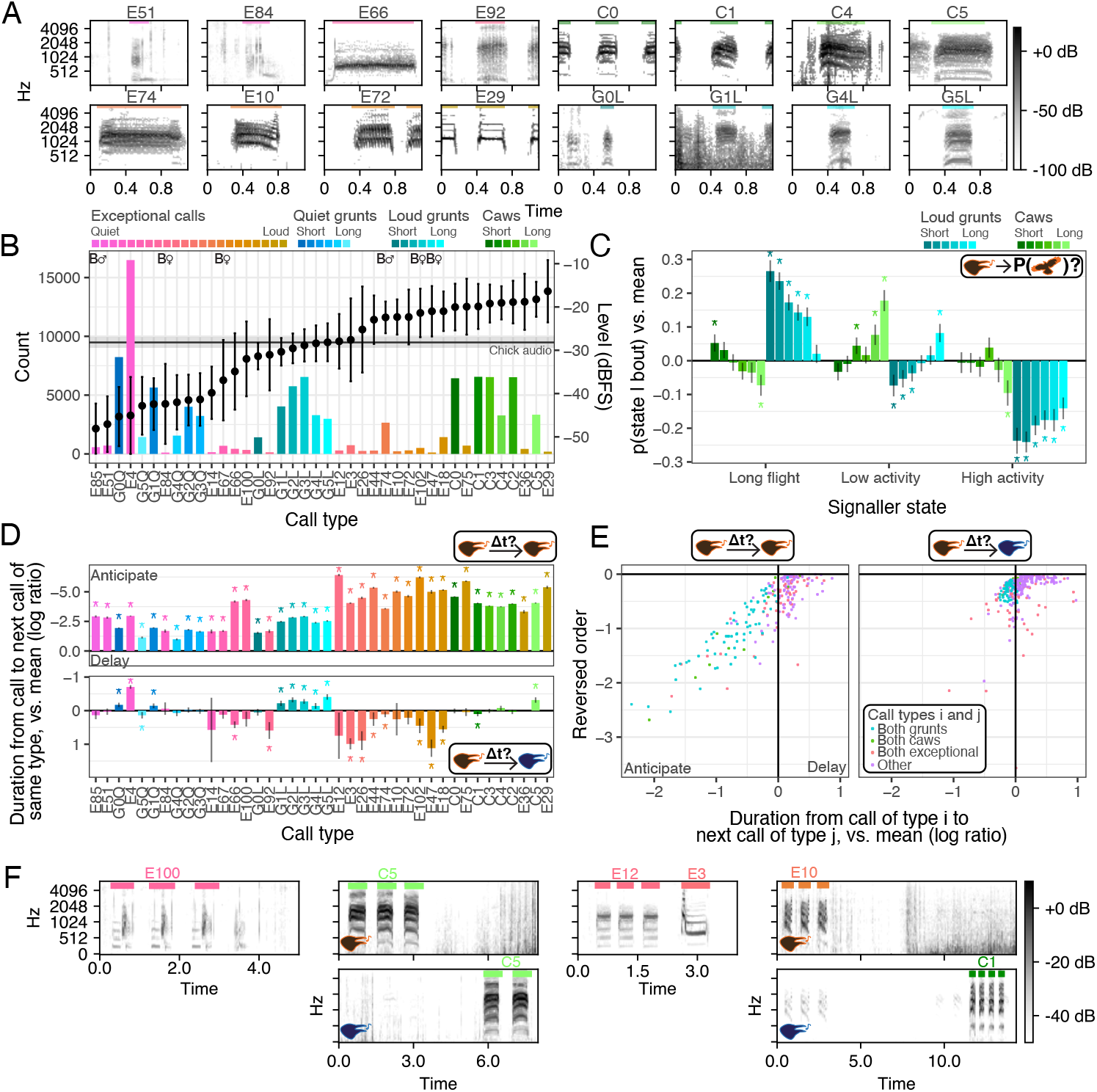
Crows have a richly structured vocal repertoire. A) Spectrograms of call types show discrete and graded variation (in grunts and caws). E: exceptional calls, GL: loud grunts, GQ: quiet grunts, C: caws. Caw and grunt integers: duration category (0=shortest, 5=longest). B) The repertoire contains rare calls, relatively quiet calls (as reference, horizontal line shows amplitude of audio containing chick vocalizations), and six social category-specific calls (B♂/B♀). Bars: count, dots/error bars: mean amplitude ±1SD. Colors: call type. C) Behaviors varied with graded acoustic variation. Both longer caws and longer grunts were more likely to indicate the low-activity state, rather than flights. D) Repetition was common in vocal sequences, while call type matching across birds was rarer. Top: All call types anticipated another call of the same type, by the same bird. Bottom: Grunts, caw5 and E4 anticipated another call of the same type, by a different bird. E) Vocal sequences (left) and exchanges (right) had pairs of calls that commonly occur together, and some of these pairs were ordered. Each dot shows a pair of call types (*v*_*i*_, *v*_*j*_). The x-axis shows the typical time from *v*_*i*_ to *v*_*j*_ (contrast to the mean call type), while the y-axis shows the reversed order. Pairs where both effects are positive (mutual delay) are not shown. F) Spectrogram examples. Column 1: Repetition in vocal sequence (D, top). Column 2: Call type matching (D, bottom). Column 3: Ordered call type pair within sequence (E, left). Column 4: Ordered call type pair in exchange (E, right). Except in B, error bars: 95% CIs; asterisks: contrast to the mean is significant with *p*_BH_ < 0.05.

The caws and grunts were graded super-categories (Figure 2A, right), accounting for, respectively, 29% and 43% of the calls. This more subtle, continuous acoustic variation could be unrelated to behavior, reflecting mainly individual variation in vocal production (Szipl, Baotic, and Kotrschal, 2026; Lee, McIvor, and Thornton, 2025): to probe this, caws and grunts were split into call types based on duration and amplitude quantiles (6 caws, labelled caw[0-5], and 12 grunts, labelled grunt[0-5]Q, grunt[0-5]L; 0 = shortest, 5 = longest, Q = quiet, L = loud). We found that behavior varied with graded acoustic features: for example, shorter caws and shorter grunts more strongly indicated the signaller was flying (Analysis 3; contrast to mean call type, *p*_BH_ < 0.05; Figure 2B).

Notably, relatively low-to intermediate-amplitude vocalizations were common. Call types with mean amplitudes lower than the mean amplitude of chick sounds, as picked up on the bird-mounted tags (grey horizontal line, Figure 2B; see Methods Section 4), accounted for 59% of vocalizations. The crows’ widely-known caws were some of their loudest vocalizations, and so give a skewed view of their communication system: our results instead suggest that communication occurred over a range of distances and was prevalent at close-range, highlighting the need for sensitive observational methods (as used here).

We also discovered structure in crow vocal sequences and exchanges by examining the duration between pairs of vocalization events (Analysis 4; Figure 2D/E). First, in agreement with previous descriptions of caws (Siriwardena, 1995), crows tended to follow their own vocalizations by another of the same type, across all call types (Analysis 4; contrast to mean call type, *p*_BH_ < 0.05; Figure 2D, top). This is easily observed in frequent repetition of calls in bouts (Figure 2F, left). In contrast, we found evidence that crows repeated the call type made by another bird only for 9 call types (Analysis 4; contrast to mean call type, *p*_BH_ < 0.05; Figure 2D, bottom). Second, we found several pairs of call types that occurred close in time, both in vocal sequences and exchanges (Figure 2E). Some of these co-occurring pairs were ordered, that is, the time between calls depended on which call type came first (see examples in Figure 2F, right). Together, these observations show that crows display previously unknown variation between call types, behaviorally relevant graded variation within call types, and sequential structure.

### 2.3 Flight calls in flocking and chick care

Vocalizations associated with flights were common and showed remarkable variation, both in their acoustic structure and behavioral associations. Overall vocalization rates were highest during the signaller’s or another bird’s flights (Analysis 2; contrast to mean activity, *p*_BH_ < 0.05; Figure 3A), and a variety of call types showed relatively high rates during flight, spanning the full range of vocalization amplitude (Analysis 2; contrast to mean activity, *p*_BH_ < 0.05; Figure 3B) and having varied associations to other behavioral states (Figure S5). This prevalence is not reflected in previous literature (Cramp, Perrins, and Brooks, 1994; Siriwardena, 1995), likely due to the difficulty of recording quiet vocalizations while the signaller is moving. Since flights coincide with transitions between different feeding sites or different activities (e.g., from foraging to feeding the chicks, resting, patrolling the territory, or mobbing), this acoustical and behavioral diversity suggests that these vocalizations are used for multiple functions related to collective movement (Dibnah et al., 2022).

**Figure 3.**
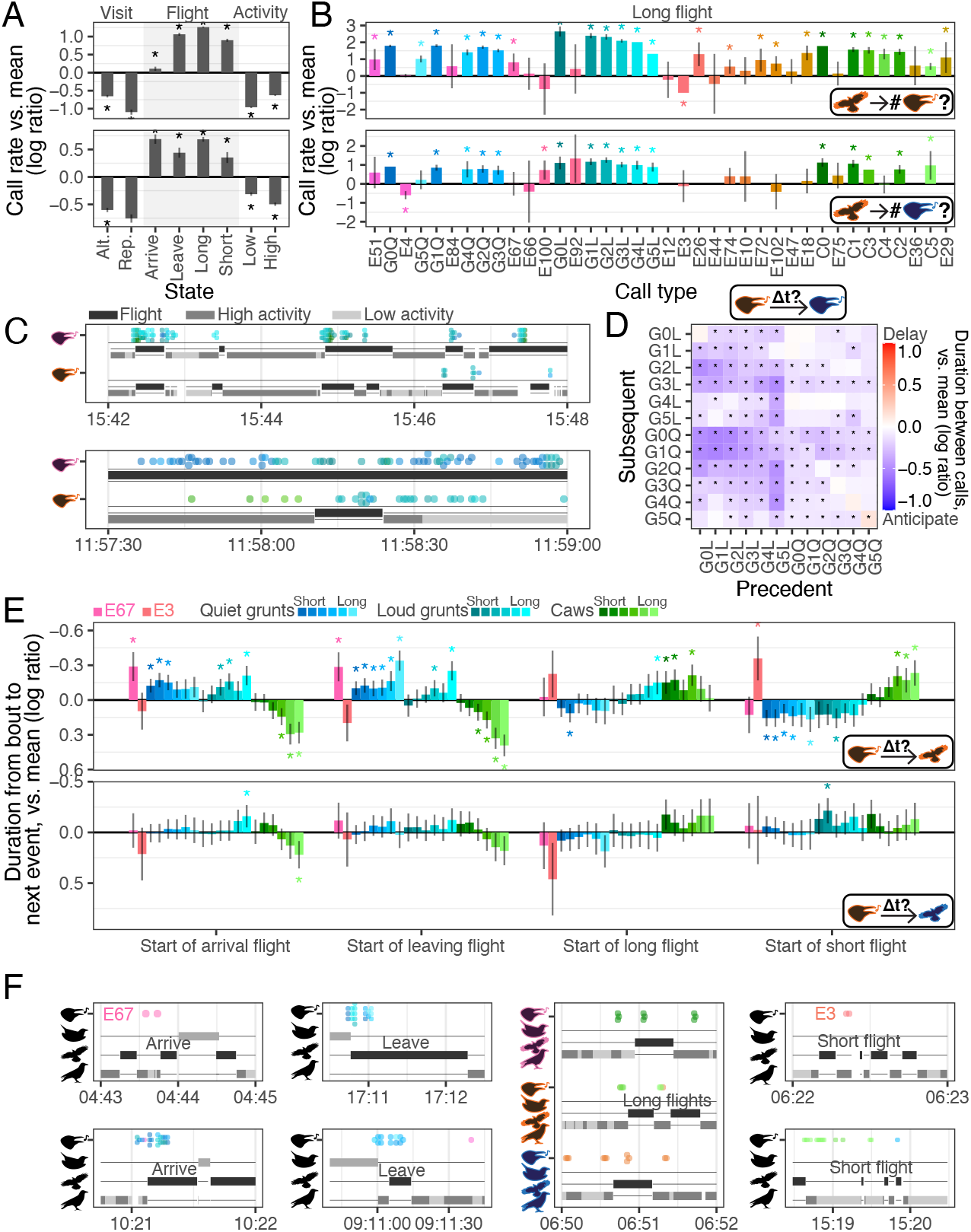
A variety of vocalizations were used during and in anticipation of flights, including to/from the nest. A) Crows had the highest overall call rate during flights (by the signaller or another bird), versus other states. B) Across multiple call types, crows called at a higher rate when they (top) or others (bottom) were making a long flight, relative to the average state. Grunts were especially common during flights. C) Sequence examples of vocalizations during flights. D) Grunts were responded to with a variety of grunts, supporting their potential function as contact calls. E) Top: Vocalizations that preceded the start of flight to nest, away from nest, long flight, and short flight. Bottom: weaker effects for preceding other’s flights. F) Sequence examples. Column 1: calls preceding start of signaller’s arrival flight. Column 2: calls preceding start of self departure flight, and ongoing during the flight. Column 3: calls preceding start of signaller’s and other’ long flight. Column 4: calls preceding start of signaller’s short flight. For all subplots, error bars: 95% CIs; asterisks: contrast to the mean is significant with *p*_BH_ < 0.05.

First, some calls appear to be used for maintaining spatial cohesion with group-mates (i.e., contact calls during movement). Grunts are the best candidate for contact calls during movement, as they tended to be made when signallers and receivers were flying (potentially simultaneously), were not social-category specific, were responded to with other grunts (Analysis 4; contrast to mean call type, *p*_BH_ < 0.05; Figure 3D), and were of low to intermediate amplitude, indicating they were employed at relatively close range. In contrast, caws were more likely to be made while the signaller was in low- or high-activity, but the receiver flying (Analysis 2; Figure S5).

Second, calls may facilitate the coordination of chick care and flocking, because they appear to announce self-movement. We found call types that preceded flights by the signaller, including flights arriving or departing at the nest (Analysis 4; contrast to mean call type, *p*_BH_ < 0.05; Figure 3E, top). Such calls could help crows coordinate chick care (Trapote et al., 2024; Mine et al., 2022), which so far had been difficult to explain mechanistically. E67 and a subset of grunts preceded both arrival and departure flights, while a subset of caws preceded other flights (long or short), indicating distinct roles for these calls. Interestingly, the breeding female was more likely to use E67, and its rate increased with chick age, suggesting that it may convey information on brood conditions (Analysis 1; pairwise contrast *p*_BH_ < 0.05; Figures S2-S3). This suggests that the female, who spends more time at the nest and has more information on chicks’ needs (Bolopo et al., 2015) (see also Figure S4), may play a distinct role in coordinating brood care. We found some weak evidence that some calls preceded others’ flights and nest visits (Figure 3E, bottom; Figure S7), warranting further investigation into whether crows use calls to direct the movement of group members. These calls may provide information beyond flights; for example, contact calls may signal group or individual identity (Keen, Meliza, and Rubenstein, 2013; Lehmann et al., 2022; Perrier et al., 2025) or calls used during flight from the nest (Figure 3A; Figure 3F, column 2) may provide information on the past nest visit, such as chick condition.

### 2.4 Adult–chick interactions during chick care

Adult–chick interactions were dominated by a single call type, E4, which may have multiple context-dependent functions. E4 was the only call with its highest rate during signaller nest visits (Analysis 2; contrast to mean behavior, *p*_BH_ < 0.05; Figure 4A). It tended to occur shortly after the signaller’s arrival at the nest (Analysis 4; contrast to mean behavior, *p*_BH_ < 0.05; Figure 4B) and before crow chick vocalizations in the signaller’s tag recording (Figure S7); E4 may thus elicit chick begging and, potentially, create opportunities for sibling negotiation (Dreiss et al., 2016). However, E4 was also used away from the nest; although rates were low, E4 increased the probability that signallers and other birds were in the low-activity state (Analysis 3; contrast to mean call type, *p*_BH_ < 0.05; Figure 4C), and crows tended to produce E4 after another adult did (Figure 2D, bottom). Therefore, a single call (or its graded variation; Figure 4F) could serve distinct functions in chick- and adult-directed contexts. Or, E4 may be generally affiliative, giving chicks an opportunity to learn group or individual signatures at the nest (Gidl et al., 2025) – a key aspect of communication in cooperative groups (Keen, Meliza, and Rubenstein, 2013; Carlson, Kelly, and Couzin, 2020).

**Figure 4.**
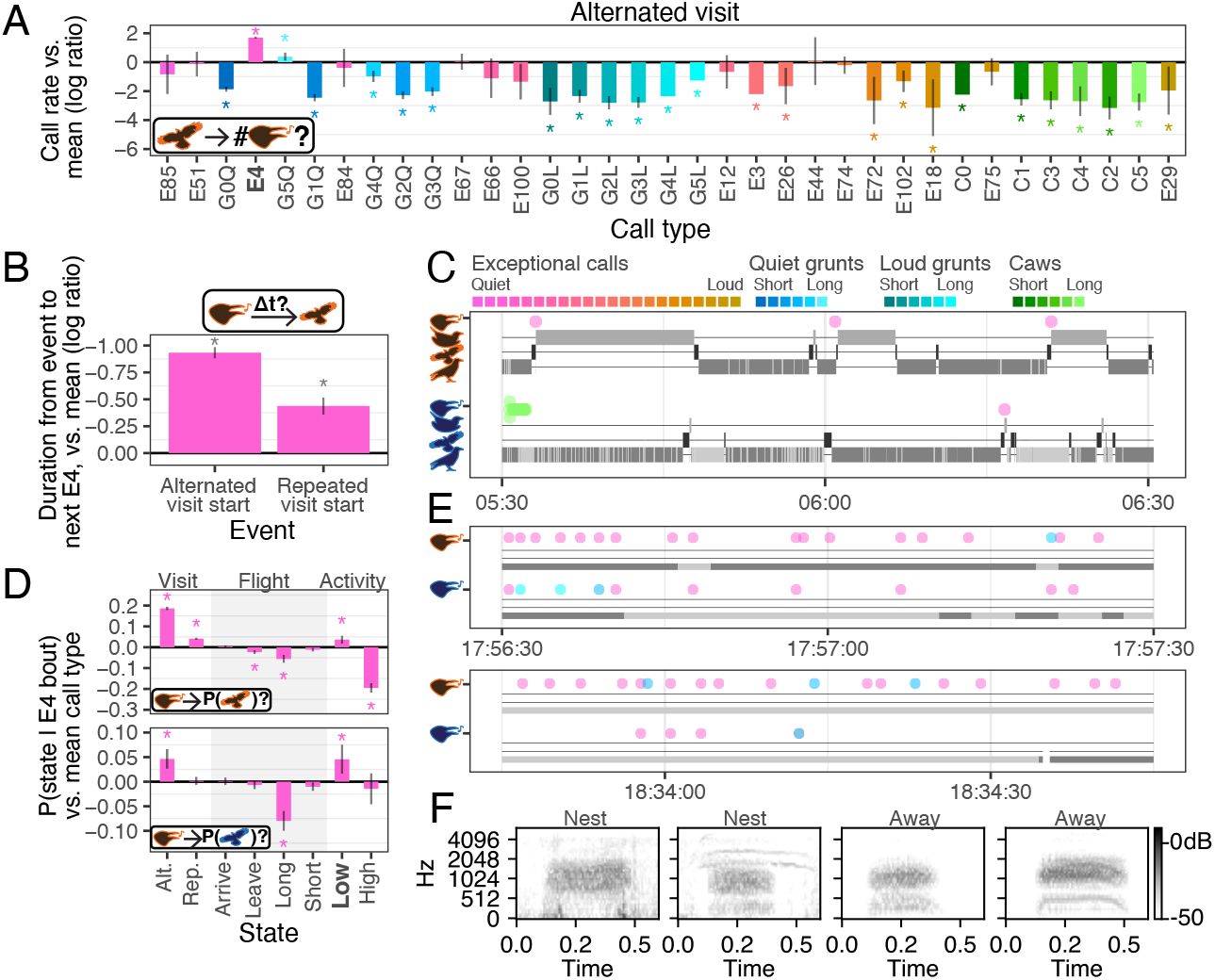
E4 can be both chick- and adult-directed. A) E4 showed a high call rate at the nest, relative to the average state. B) The beginning of a signaller’s nest visits anticipated E4: the duration between these events and E4 was small, relative to the mean over precedent events. C) A sequence example showing E4 (pink) produced at the beginning of four nest visits, which occur within an hour. D) E4 also occured away from the nest. In particular, E4 increased the probability that signaller and other birds are in the low-activity state. Bars: probability of being in the specified behavioral state, given the occurrence of an E4 bout, relative to the mean call type. E) Two sequence examples showing birds producing E4 while they were in low- and high-activity states, rather than at the nest. F) Spectrogram examples of E4 at and away from the nest. For all subplots, error bars: 95% CIs; asterisks: contrast to the mean is significant with *p*_BH_ < 0.05.)

### 2.5 Territorial display

We also identified a system of vocalizations that likely play a role in marking territorial boundaries. Territorial calls are expected to be relatively loud, since they are directed to crows outside the group, and to be especially frequent at dawn (Cramp, Perrins, and Brooks, 1994). Also, we would expect group members to chorus, that is, use the same type of vocalization to reinforce the signal directed to outsiders (Bradbury, Vehrencamp, et al., 1998). We found that out of 20 call types with amplitudes greater than or equal to mean chick vocalization amplitude (Figure 2B), all caw calls and three exceptional calls (E74, E75 and E3) showed a high rate of usage in the morning, and subsided in the central hours of the day (Analysis 1; contrast to the mean over hours, *p*_BH_ < 0.05; Figure 5A), supporting a territorial function. All these adult calls increased in rate with chick age (Analysis 1; *p*_BH_ < 0.05; Figure 5E, right), potentially signalling greater motivation to defend a high-value territory, although mixed results were found for the number of chicks (Figure 5E, left). These calls varied from each other across several other factors: the exceptional calls were rarer (Figure 5A), were used during different activities of the signaller and other birds (Analyses 2-3; contrast to the mean, *p*_BH_ < 0.05; Figure 5B), and E3 and caws preceded flights (Figure 5E), while the others did not. The rate of all vocalizations except E74 did not vary with social category, suggesting that all adult group members participate in territorial advertising, not just the dominant breeders. It was breeding males who preferentially used E74 (Analysis 1; pairwise contrast *p*_BH_ < 0.05; Figure 2B) and who also spent the most time flying (Figure S4), suggesting they may patrol more and may contribute to coordinating territorial displays (a situation resembling the breeding females’ use of E67 in chick care).

**Figure 5.**
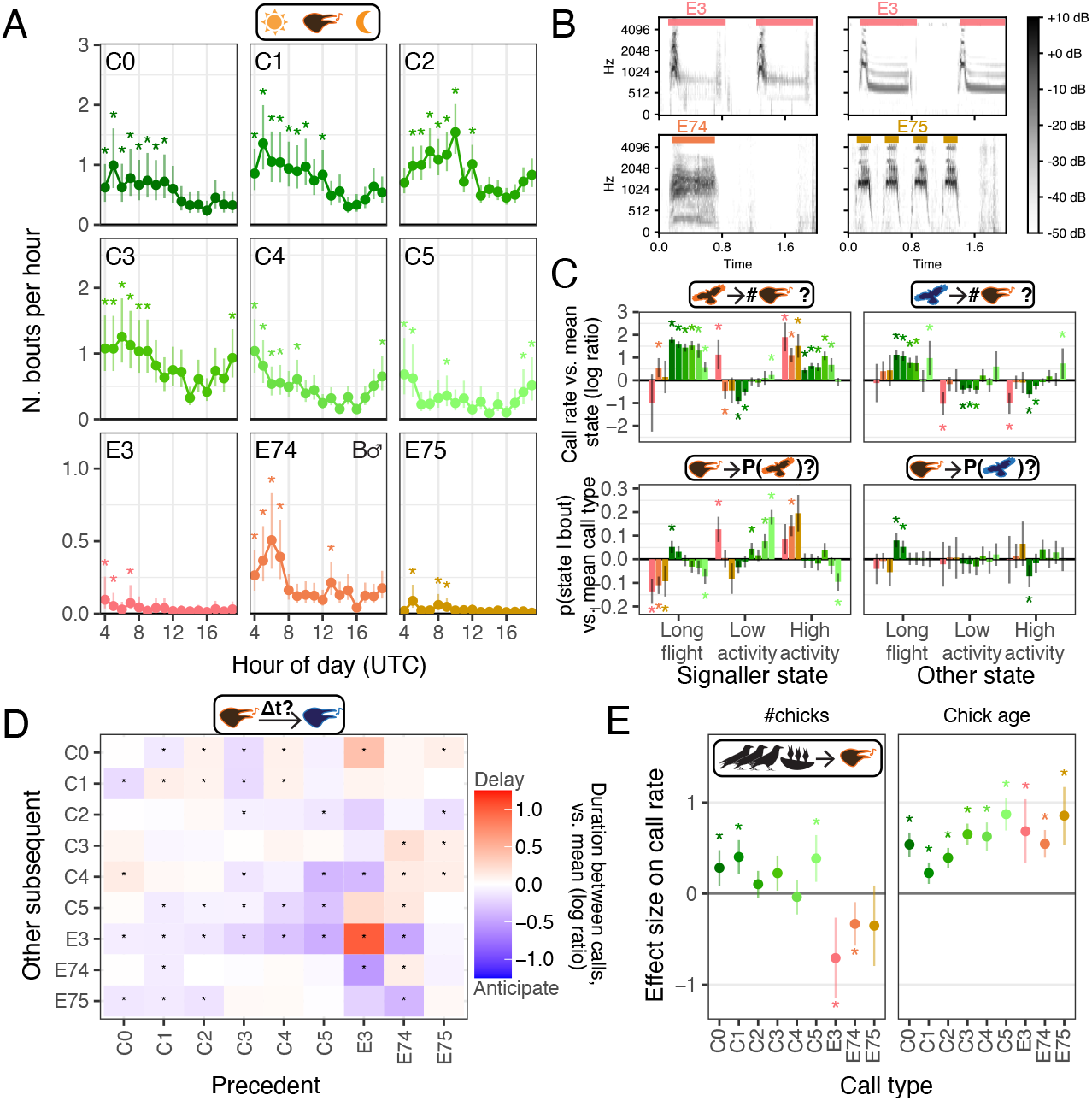
A variety of loud vocalizations are used in the morning, consistent with territorial display. A) All caw calls and three exceptional calls (E74, E75 and E3) showed a high rate of emission early in the morning relative to other hours of day, and subsided in the central hours of the day, supporting a territorial function. The calls vary in their overall rate (the exceptional calls are rarer) and their temporal profile (C2 peaked in the middle of the day, while longer-duration caws had an additional evening peak). E74 differs in that breeding males tend to produce it at higher rates. Asterisks: contrast to the mean is significant with *p*_BH_ < 0.05, for positive effects only. B) The exceptional call types are acoustically diverse. C) The calls vary with respect to behavioural states, both of the signaller and other birds. Asterisks: contrast to the mean is significant with *p*_BH_ < 0.05. D) For the selected calls, crows do not tend to chorus (i.e., do not respond with the same call type) but rather respond with a different type. Asterisks: contrast to the mean is significant with *p*_BH_ < 0.05. E) The selected calls vary in how their rate is related to group composition, specifically the number (left) and the age of the chicks (right). The rate of all call types increased with the age of the chicks, while the number of chicks had a varied effect across call types. Asterisks: effects are significant with *p*_BH_ < 0.05. For all subplots, error bars: 95% CIs.

In contrast to our expectations, we found mixed evidence for chorusing behavior using these call types (Demartsev, Averly, et al., 2024). For example, while caws were often followed by other caws, these repeated caws were typically of different duration, and caws were also often followed by exceptional call types (Analysis 4; contrasts to mean call type, *p*_BH_ < 0.05; Figure 5E). Beyond simply marking the territory, this diversity of vocalizations may serve different functions consistent with their varied behavioral associations (Savagian and Riehl, 2023), or advertise group composition (Hall and Magrath, 2007) – a potentially crucial signal in social or competitive interactions. Another deviation from our expectations regarding territorial display was that longer-duration caws were common in the evening hours. The graded variation may be associated not only with morning territorial displays, but also with roosting behavior: the longest caws also tended to indicate the signaller’s low-activity state (Figure 2C), which is likely to encompass perching, and tended to precede short rather than long flights (Figure 3E), suggesting fewer long-distance transitions.

### 2.6 Other exceptional calls

Some exceptional calls were not involved in flocking, chick care, or territorial display. While vocalization rates were lower overall during the high- and low-activity states (Figure 3A), a large variety of rare exceptional calls occurred while signallers were in these states (Figure S5). Possible behaviors during these states include foraging and roosting, which crows are known to do cohesively (Baglione, Canestrari, Marcos, Griesser, et al., 2002), as well as food caching, sentineling, bathing, and resting. More refined behavioral characterization and experimentally manipulated contexts (Rutz, Bronstein, et al., 2023; Bugnyar, Kijne, and Kotrschal, 2001) could help distinguish these calls, and may also provide further insight into flight-related calls, as they occur during behavioral transitions.

## 3 Concluding remarks

Integrating results across the multiscale analyses produced a step-change in our understanding of crow communication. By using large-scale, fine-timescale observational data, we were able to map functionally diverse sets of calls in three different joint behaviors: flocking, chick care, and territorial display. Hypothesized functions for call types were informed by multiple types of associations, patterns that emerged across contexts (e.g., away vs. at the nest), and in consideration of other call types (e.g., vocal exchange structure for caws vs. grunts), suggesting the importance of treating the repertoire as a communication system rather than its components in isolation. While our findings are correlational in nature, they provide critical insights into communication, including aspects such as self-movement announcement that are beyond the reach of experimental manipulation (Salis, Martin, and Girard-Buttoz, 2025).

We uncovered that crows use a rich vocal repertoire, which seems commensurate with the social complexity inherent in these birds’ lives. Comparative studies that explore the relationship between social and vocal complexity typically ignore potential dependencies between the functions of a vocalization and its social contexts, which may undermine conclusions about the causes of signal variation (Freeberg, Dunbar, and Ord, 2012). Repertoire-wide mapping could form the basis for fine-grained comparative studies on function, for example, with the same crow population at different times of year when social interactions vary (e.g., courtship, (Gill et al., 2015); winter roosts with extra-group members, (Gómez et al., 2024; Boucherie et al., 2019)), with populations of carrion crows that do not breed cooperatively (Baglione, Marcos, and Canestrari, 2002), and with other corvids, who live in diverse societies (Clayton and Emery, 2007).

## Contributions

Conceptualization: MC, BH, JL, CR, EC, VB, DC Methodology: MC, BH, CG, AM, MB, SCK, CR, EC, VB, DC Software: MC, BH, JL, LM, MM, DR Formal analysis: MC, BH Investigation: MC, BH, MV, ET, VB, DC Resources: MA, VB, DC, CR, MV Data curation: MC, BH, CG, AM, MB, JL, VB, DC Writing – original draft: MC, BH, VB, DC Writing – review & editing: MC, BH, CG, AM, MB, LSJ, SCK, LM, MV, DEB, CR, EC, VB, DC Visualization: MC, LSJ, EC Supervision: MC, BH, EC, MG, OP, VB, DC, CR Project administration: MC, BH, EC, OP, VB, DC, CR Funding acquisition: BH, OP, CR, VB, DC

## Acknowledgements

We are grateful to the crows in this study. We acknowledge Mark Peter Johnson for his contributions to sensor design, data collection, and methods review. We acknowledge the staff of the Earth Species Project for help with logistics, fundraising, and feedback on the project, including Masato Hagiwara, Felix Effenberger, Gagan Narula, and Diane Kim for constructive discussions, and Ellen Gilsenan-McMahon, Laura Hay, and Sylvie Manning for organizational support. We thank Phoebe Koenig for input on analysis design, and Luke Hewitt for input on analysis design and manuscript review. Earth Species Project also acknowledges our core supporters.

Data collection was funded by the Spanish Ministerio de Ciencia e Innovación (MCIN/AEI/10.13039/ 501100011033) to VB. Data analyses and writing were covered by the Project PID2024-159215NB-100, Plan Nacional de I+D+i, Ministerio de Ciencia e Innovación y Agencia Estatal de Investigación, to DC. CR acknowledges funding from the Biotechnology and Biological Sciences Research Council (BB/S018484/1). MV acknowledges funding from Xunta de Galicia (Spain) (Grant ED431B 2024/23). DEB acknowledges funding from Schmidt Sciences. This project was supported (in part) by a grant from the National Geographic Society to Earth Species Project.

## References

Baglione, Vittorio, Daniela Canestrari, Elisa Chiarati, Ruben Vera, and José M Marcos (2010). “Lazy group members are substitute helpers in carrion crows.” In: Proceedings of the Royal Society B: Biological Sciences 277.1698, pp. 3275–3282.

Baglione, Vittorio, Daniela Canestrari, Maddie Cusimano, Benjamin Hoffman, Victor Moreno, and Eva Trapote (2025). “Capturing vocal communication in a free-living corvid: high-resolution data from low-impact miniaturized tags.” In: Animal Cognition 28.1, p. 85.

Baglione, Vittorio, Daniela Canestrari, José M Marcos, and Jan Ekman (2003). “Kin selection in cooperative alliances of carrion crows.” In: Science 300.5627, pp. 1947–1949.

Baglione, Vittorio, Daniela Canestrari, José M Marcos, and Jan Ekman (2006). “Experimentally increased food resources in the natal territory promote offspring philopatry and helping in cooperatively breeding carrion crows.” In: Proceedings of the Royal Society B: Biological Sciences 273.1593, pp. 1529–1535.

Baglione, Vittorio, Daniela Canestrari, José M Marcos, Michael Griesser, and Jan Ekman (2002). “History, environment and social behaviour: experimentally induced cooperative breeding in the carrion crow.” In: Proceedings of the Royal Society of London. Series B: Biological Sciences 269.1497, pp. 1247–1251.

Baglione, Vittorio, José M Marcos, and Daniela Canestrari (2002). “Cooperatively breeding groups of carrion crow (Corvus corone corone) in northern Spain.” In: The Auk 119.3, pp. 790–799.

Baglione, Vittorio, José M Marcos, Daniela Canestrari, and Jan Ekman (2002). “Direct fitness benefits of group living in a complex cooperative society of carrion crows, Corvus corone corone.” In: Animal Behaviour 64.6, pp. 887–893.

Barker, Alison J, Grigorii Veviurko, Nigel C Bennett, Daniel W Hart, Lina Mograby, and Gary R Lewin (2021). “Cultural transmission of vocal dialect in the naked mole-rat.” In: Science 371.6528, pp. 503–507.

Bates, Douglas, Martin Mächler, Ben Bolker, and Steve Walker (2015). “Fitting Linear Mixed-Effects Models Using lme4.” In: Journal of Statistical Software 67.1, pp. 1–48. DOI: 10.18637/jss.v067.i01.

Bolopo, Diana, Daniela Canestrari, José M Marcos, and Vittorio Baglione (2015). “Nest sanitation in cooperatively breeding Carrion Crows.” In: The Auk: Ornithological Advances 132.3, pp. 604–612.

Boucherie, Palmyre H, Matthias-Claudio Loretto, Jorg JM Massen, and Thomas Bugnyar (2019). “What constitutes “social complexity” and “social intelligence” in birds? Lessons from ravens.” In: Behavioral Ecology and Sociobiology 73.1, p. 12.

Bradbury, Jack W, Sandra Lee Vehrencamp, et al. (1998). Principles of animal communication. Vol. 132. Sinauer Associates Sunderland, MA.

Bugnyar, Thomas, Maartje Kijne, and Kurt Kotrschal (2001). “Food calling in ravens: are yells referential signals?” In: Animal Behaviour 61.5, pp. 949–958.

Caffrey, Carolee (2000). “Marking crows.” In: North American Bird Bander 26.4, p. 2.

Cakir, Emre, Toni Heittola, Heikki Huttunen, and Tuomas Virtanen (2015). “Polyphonic sound event detection using multi label deep neural networks.” In: 2015 International Joint Conference on Neural Networks (IJCNN). IEEE, pp. 1–7.

Canestrari, Daniela, Diana Bolopo, Ted CJ Turlings, Gregory Röder, José M Marcos, and Vittorio Baglione (2014). “From parasitism to mutualism: unexpected interactions between a cuckoo and its host.” In: Science 343.6177, pp. 1350–1352.

Canestrari, Daniela, José M Marcos, and Vittorio Baglione (2005). “Effect of parentage and relatedness on the individual contribution to cooperative chick care in carrion crows Corvus corone corone.” In: Behavioral Ecology and Sociobiology 57.5, pp. 422–428.

Canestrari, Daniela, José M Marcos, and Vittorio Baglione (2009). “Cooperative breeding in carrion crows reduces the rate of brood parasitism by great spotted cuckoos.” In: Animal Behaviour 77.5, pp. 1337–1344.

Carlson, Nora V, E McKenna Kelly, and Iain Couzin (2020). “Individual vocal recognition across taxa: a review of the literature and a look into the future.” In: Philosophical Transactions of the Royal Society B: Biological Sciences 375.1802, p. 20190479.

Chen, Sanyuan, Yu Wu, Chengyi Wang, Shujie Liu, Daniel Tompkins, Zhuo Chen, and Furu Wei (2022). “Beats: Audio pre-training with acoustic tokenizers.” In: arXiv preprint arXiv:2212.09058.

Chiarati, Elisa, Daniela Canestrari, Rubén Vera, José M Marcos, and Vittorio Baglione (2010). “Linear and stable dominance hierarchies in cooperative carrion crows.” In: Ethology 116.4, pp. 346–356.

Clayton, Nicola S and Nathan J Emery (2007). “The social life of corvids.” In: Current Biology 17.16, R652–R656.

Committee, ASAB Ethical Committee/ABS Animal Care et al. (2024). “Guidelines for the ethical treatment of nonhuman animals in behavioural research and teaching.” In: Animal Behaviour 207, pp. I–XI.

Cramp, S, CM Perrins, and DJ Brooks (1994). Handbook of the birds of Europe, the Middle East and North Africa: the birds of the Western Palearctic. Vol. VIII: Crows to finches. Oxford: Oxford University Press.

Demartsev, Vlad, Baptiste Averly, Lily Johnson-Ulrich, Vivek H Sridhar, Leonardos Leonardos, Alexander Q Vining, Mara Thomas, Marta B Manser, and Ariana Strandburg-Peshkin (2024). “Mapping vocal interactions in space and time differentiates signal broadcast versus signal exchange in meerkat groups.” In: Philosophical Transactions of the Royal Society B: Biological Sciences 379.1905, p. 20230188.

Demartsev, Vlad, Andrew S Gersick, Frants H Jensen, Mara Thomas, Marie A Roch, Marta B Manser, and Ariana Strandburg-Peshkin (2023). “Signalling in groups: New tools for the integration of animal communication and collective movement.” In: Methods in Ecology and Evolution 14.8, pp. 1852–1863.

Denton, Tom, Scott Wisdom, and John R Hershey (2022). “Improving bird classification with unsupervised sound separation.” In: ICASSP 2022-2022 IEEE International Conference on Acoustics, Speech and Signal Processing (ICASSP). IEEE, pp. 636–640.

Dibnah, Alex J, James E Herbert-Read, Neeltje J Boogert, Guillam E McIvor, Jolle W Jolles, and Alex Thornton (2022). “Vocally mediated consensus decisions govern mass departures from jackdaw roosts.” In: Current Biology 32.10, R455–R456.

Dreiss, Amélie N, Florence Gaime, Alice Delarbre, Letizia Moroni, Mélissa Lenarth, and Alexandre Roulin (2016). “Vocal communication regulates sibling competition over food stock.” In: Behavioral Ecology and Sociobiology 70.6, pp. 927–937.

Eleuteri, Vesta, Lucy Bates, Jake Rendle-Worthington, Catherine Hobaiter, and Angela Stoeger (2024). “Multimodal communication and audience directedness in the greeting behaviour of semi-captive African savannah elephants.” In: Communications Biology 7.1, p. 472.

Elie, Julie E and Frédéric E Theunissen (2018). “Zebra finches identify individuals using vocal signatures unique to each call type.” In: Nature communications 9.1, p. 4026.

Ellegren, Hans and Anna-Karin Fridolfsson (1997). “Male–driven evolution of DNA sequences in birds.” In: Nature Genetics 17.2, pp. 182–184.

Elliott, Kyle H (2025). “Sweat the small stuff: A review of the use of accelerometers to estimate energy expenditure in wild animals.” In: Journal of Animal Ecology 94.12, pp. 2362–2375.

Fleischer, Robert C, William I Boarman, Elena G Gonzalez, Alvaro Godinez, Kevin E Omland, Sarah Young, Lauren Helgen, Gracia Syed, and Carl E Mcintosh (2008). “As the raven flies: using genetic data to infer the history of invasive common raven (Corvus corax) populations in the Mojave Desert.” In: Molecular Ecology 17.1, pp. 464–474.

Freeberg, Todd M, Robin IM Dunbar, and Terry J Ord (2012). “Social complexity as a proximate and ultimate factor in communicative complexity.” In: Philosophical Transactions of the Royal Society B: Biological Sciences 367.1597, pp. 1785–1801.

Gidl, Hannah, Sara Binder, Anna N Osiecka, and Barbara C Klump (2025). “The ontogeny of vocal identity in carrion crows (Corvus corone).” In: Animal Cognition.

Gill, Lisa F, Wolfgang Goymann, Andries Ter Maat, and Manfred Gahr (2015). “Patterns of call communication between group-housed zebra finches change during the breeding cycle.” In: eLife 4, e07770.

Gómez, Rubén Vera, Vittorio Baglione, Elisa Chiarati, and Daniela Canestrari (2024). “Social preference persists at roosting aggregations in a cooperatively breeding bird.” In: Animal Behaviour 218, pp. 87–94.

Gomila, Robin (2021). “Logistic or linear? Estimating causal effects of experimental treatments on binary outcomes using regression analysis.” In: Journal of Experimental Psychology: General 150.4, p. 700.

Gould, James (1974). “Honey bee communication.” In: Nature 252, pp. 300–301. DOI: 10.1038/252300a0.

Hagiwara, Masato (2023). “Aves: Animal vocalization encoder based on self-supervision.” In: ICASSP 2023-2023 IEEE International Conference on Acoustics, Speech and Signal Processing (ICASSP). IEEE, pp. 1–5.

Hall, Michelle L and Robert D Magrath (2007). “Temporal coordination signals coalition quality.” In: Current Biology 17.11, R406–R407.

Hellevik, Ottar (2009). “Linear versus logistic regression when the dependent variable is a dichotomy.” In: Quality & quantity 43.1, pp. 59–74.

Hoffman, Benjamin, Maddie Cusimano, Vittorio Baglione, Daniela Canestrari, Damien Chevallier, Dominic L DeSantis, Lorène Jeantet, Monique A Ladds, Takuya Maekawa, Vicente Mata-Silva, et al. (2024). “A benchmark for computational analysis of animal behavior, using animal-borne tags.” In: Movement Ecology 12.1, p. 78.

Jones, Owen R and Jinliang Wang (2010). “COLONY: a program for parentage and sibship inference from multilocus genotype data.” In: Molecular Ecology Resources 10.3, pp. 551–555.

Kalinowski, Steven T, Mark L Taper, and Tristan C Marshall (2007). “Revising how the computer program CERVUS accommodates genotyping error increases success in paternity assignment.” In: Molecular Ecology 16.5, pp. 1099–1106.

Keen, Sara C, C Daniel Meliza, and Dustin R Rubenstein (2013). “Flight calls signal group and individual identity but not kinship in a cooperatively breeding bird.” In: Behavioral Ecology 24.6, pp. 1279–1285.

Kelso, Erin C and Paul A Verrell (2002). “Do male veiled chameleons, Chamaeleo calyptratus, adjust their courtship displays in response to female reproductive status?” In: Ethology 108.6, pp. 495–512.

Kingma, Diederik P and Jimmy Ba (2014). “Adam: A method for stochastic optimization.” In: arXiv preprint arXiv:1412.6980.

Kondo, Noriko and Shigeru Watanabe (2009). “Contact calls: information and social function.” In: Japanese Psychological Research 51.3, pp. 197–208.

Kuznetsova, Alexandra, Per B. Brockhoff, and Rune H. B. Christensen (2017). “lmerTest Package: Tests in Linear Mixed Effects Models.” In: Journalof Statistical Software 82.13, pp. 1–26. DOI: 10.18637/jss.v082.i13.

Lee, Victoria E, Guillam E McIvor, and Alex Thornton (2025). “Wild jackdaws recognise the contact calls of their mate.” In: Animal Cognition 28.1, p. 97.

Lehmann, Kenna DS, Frants H Jensen, Andrew S Gersick, Ariana Strandburg-Peshkin, and Kay E Holekamp (2022). “Long-distance vocalizations of spotted hyenas contain individual, but not group, signatures.” In: Proceedings of the Royal Society B: Biological Sciences 289.1979.

Leighton, Gavin M (2017). “Cooperative breeding influences the number and type of vocalizations in avian lineages.” In: Proceedings of the Royal Society B: Biological Sciences 284.1868, p. 20171508.

Lenth, Russell V. and Julia Piaskowski (2026). emmeans: Estimated Marginal Means, aka Least-Squares Means. R package version 2.0.2. URL: https://rvlenth.github.io/emmeans/.

Li, Shou-Hsien, Yi-Jiun Huang, and JL Brown (1997). “Isolation of tetranucleotide microsatellites from the Mexican jay Aphelocoma ultramarina.” In: Molecular Ecology 6.5, pp. 499–501.

Liao, Diana A, Katharina F Brecht, Lena Veit, and Andreas Nieder (2024). “Crows “count” the number of self-generated vocalizations.” In: Science 384.6698, pp. 874–877.

Magrath, Robert D, Tonya M Haff, Pamela M Fallow, and Andrew N Radford (2015). “Eavesdropping on heterospecific alarm calls: from mechanisms to consequences.” In: Biological Reviews 90.2, pp. 560–586.

Mahon, Louis, Benjamin Hoffman, Logan James, Maddie Cusimano, Masato Hagiwara, Sarah C Woolley, Felix Effenberger, Sara Keen, Jen-Yu Liu, and Olivier Pietquin (2025). “Robust detection of overlapping bioacoustic sound events.” In: arXiv preprint arXiv:2503.02389.

Marzluff, John and Tony Angell (2013). Gifts of the crow: how perception, emotion, and thought allow smart birds to behave like humans. Simon and Schuster.

Mas, Flore and Mathias Kölliker (2008). “Maternal care and offspring begging in social insects: chemical signalling, hormonal regulation and evolution.” In: Animal Behaviour 76.4, pp. 1121–1131.

McGowan, Kevin J and Glen E Woolfenden (1989). “A sentinel system in the Florida scrub jay.” In: Animal Behaviour 37, pp. 1000–1006.

Meirmans, Patrick G (2020). “Genodive version 3.0: Easy-to-use software for the analysis of genetic data of diploids and polyploids.” In: Molecular Ecology Resources 20.4, pp. 1126–1131.

Merino Recalde, Nilo (2023). “pykanto: A python library to accelerate research on wild bird song.” In: Methods in Ecology and Evolution 14.8, pp. 1994–2002.

Mine, Joseph G, Katie E Slocombe, Erik P Willems, Ian C Gilby, Miranda Yu, Melissa Emery Thompson, Martin N Muller, Richard W Wrangham, Simon W Townsend, and Zarin P Machanda (2022). “Vocal signals facilitate cooperative hunting in wild chimpanzees.” In: Science advances 8.30, eabo5553.

Nathan, Ran, Orr Spiegel, Scott Fortmann-Roe, Roi Harel, Martin Wikelski, and Wayne M Getz (2012). “Using tri-axial acceleration data to identify behavioral modes of free-ranging animals: general concepts and tools illustrated for griffon vultures.” In: Journal of Experimental Biology 215.6, pp. 986–996.

Perrier, Léo, Lény Lego, Tristan Cladière, Martin Blanchard, Lindelani Makuya, Wiebke Berns, Aurélie Pradeau, Carsten Schradin, Michael D Greenfield, Nicolas Mathevon, et al. (2025). “Ultrasonic signals support a large-scale communication landscape in wild mice.” In: Current Biology 35.19, pp. 4837–4844.

Price, Tabitha, Philip Wadewitz, Dorothy Cheney, Robert Seyfarth, Kurt Hammerschmidt, and Julia Fischer (2015). “Vervets revisited: A quantitative analysis of alarm call structure and context specificity.” In: Scientific reports 5.1, p. 13220.

Rousset, Francois (2008). “genepop’007: a complete re-implementation of the genepop software for Windows and Linux.” In: Molecular Ecology Resources 8.1, pp. 103–106.

Ruiter, Stacy L. de, Mark Johnson, K. Alex Shorter, Lucía Martina Martín López, and Joaquin T. Gabaldon (2020). Biologging Tools Project. URL: http://animaltags.org/.

Rutz, Christian, Michael Bronstein, Aza Raskin, Sonja C Vernes, Katherine Zacarian, and Damián E Blasi (2023). “Using machine learning to decode animal communication.” In: Science 381.6654, pp. 152–155.

Rutz, Christian and Jolyon Troscianko (2013). “Programmable, miniature video-loggers for deployment on wild birds and other wildlife.” In: Methods in Ecology and Evolution 4.2, pp. 114–122.

Salis, Ambre, Killian Martin, and Cédric Girard-Buttoz (2025). “Challenges and new opportunities in deciphering the meaning of corvid call sequences.” In: Animal Cognition 28.1, p. 95.

Savagian, Amanda and Christina Riehl (2023). “Group chorusing as an intragroup signal in the greater ani, a communally breeding bird.” In: Ethology 129.2, pp. 63–73.

Siriwardena, Gavin Mark (1995). Aspects of vocal communication in the carrion crow, Corvus corone corone. University of Leicester (United Kingdom).

Smeele, Simeon Q, Stephen A Tyndel, Barbara C Klump, Gustavo Alarcón-Nieto, and Lucy M Aplin (2024). “callsync: An R package for alignment and analysis of multi-microphone animal recordings.” In: Ecology and Evolution 14.5, e11384.

Stidsholt, Laura, Mark Johnson, Kristian Beedholm, Lasse Jakobsen, Kathrin Kugler, Signe Brinkløv, Angeles Salles, Cynthia F Moss, and Peter Teglberg Madsen (2019). “A 2.6-g sound and movement tag for studying the acoustic scene and kinematics of echolocating bats.” In: Methods in Ecology and Evolution 10.1, pp. 48–58.

Stowell, Dan (2022). “Computational bioacoustics with deep learning: a review and roadmap.” In: PeerJ 10, e13152.

Stowell, Dan, Emmanouil Benetos, and Lisa F Gill (2017). “On-bird sound recordings: automatic acoustic recognition of activities and contexts.” In: IEEE/ACMTransactionson Audio, Speech, and Language Processing 25.6, pp. 1193–1206.

Szipl, Georgine, Anton Baotic, and Kurt Kotrschal (2026). “Variability and individuality in the contact calls of jackdaws (Corvus monedula).” In: Animal Cognition 29.1, p. 2.

Thomas, Mara, Frants H Jensen, Baptiste Averly, Vlad Demartsev, Marta B Manser, Tim Sainburg, Marie A Roch, and Ariana Strandburg-Peshkin (2022). “A practical guide for generating unsupervised, spectrogram-based latent space representations of animal vocalizations.” In: Journal of Animal Ecology 91.8, pp. 1567–1581.

Tibbetts, Elizabeth A, Juanita Pardo-Sanchez, and Chloe Weise (2022). “The establishment and maintenance of dominance hierarchies.” In: Philosophical Transactionsofthe Royal Society B: Biological Sciences 377.1845.

Trapote, Eva, Víctor Moreno-González, Daniela Canestrari, Christian Rutz, and Vittorio Baglione (2024). “Fitness benefits of alternated chick provisioning in cooperatively breeding carrion crows.” In: Journal of Animal Ecology 93.1, pp. 95–108.

Tuia, Devis, Benjamin Kellenberger, Sara Beery, Blair R Costelloe, Silvia Zuffi, Benjamin Risse, Alexander Mathis, Mackenzie W Mathis, Frank Van Langevelde, Tilo Burghardt, et al. (2022). “Perspectives in machine learning for wildlife conservation.” In: Nature communications 13.1, p. 792.

Waas, Joseph R (1991). “The risks and benefits of signalling aggressive motivation: a study of cave-dwelling little blue penguins.” In: Behavioral Ecology and Sociobiology 29.2, pp. 139–146.

Wang, Ning, Jianqiang LI, Yingying Liu, and Zhengwang Zhang (2010). “Improvement on molecular sex identification primers for Passeriform bird species.” In: Avian Research 1.1, pp. 65–69. ISSN: 2055-6187 (Print). DOI: 10.5122/cbirds.2009.0009.

Wascher, Claudia AF and Sam Reynolds (2025). “Vocal communication in corvids: a systematic review.” In: Animal Behaviour 221, p. 123073.

Webster, Michael M and Christian Rutz (2020). “How STRANGE are your study animals?” In:Nature 582.7812, pp. 337–340.

Wild, Timm A, Georg Wilbs, Dina KN Dechmann, Jenna E Kohles, Nils Linek, Sierra Mattingly, Nina Richter, Spyros Sfenthourakis, Haris Nicolaou, Elena Erotokritou, et al. (2024). “Time synchronisation for millisecond-precision on bio-loggers.” In: Movement Ecology 12.1, p. 71.

Wilson, Rory P, Luca Börger, Mark D Holton, D Michael Scantlebury, Agustina Gómez-Laich, Flavio Quintana, Frank Rosell, Patricia M Graf, Hannah Williams, Richard Gunner, et al. (2020). “Estimates for energy expenditure in free-living animals using acceleration proxies: A reappraisal.” In: Journal of Animal Ecology 89.1, pp. 161–172.

Wilson, Rory P, Craig R White, Flavio Quintana, Lewis G Halsey, Nikolai Liebsch, Graham R Martin, and Patrick J Butler (2006). “Moving towards acceleration for estimates of activity-specific metabolic rate in free-living animals: the case of the cormorant.” In: Journal of Animal Ecology 75.5, pp. 1081–1090.

Zeh, Julia M, Valeria Perez-Marrufo, Dana L Adcock, Frants H Jensen, Kaitlyn J Knapp, Jooke Robbins, Jennifer E Tackaberry, Mason Weinrich, Ari S Friedlaender, David N Wiley, et al. (2024). “Caller identification and characterization of individual humpback whale acoustic behaviour.” In: Royal Society Open Science 11.3.

