## Supplementary Materials for "Repertoire-behavior mapping reveals signal functions in cooperatively breeding crows"

### Materials and Methods

#### *1 Study area and population*

The study area is located in a rural region of northern Spain (42°37'N, 5°26'W) where long-term research on a population of carrion crows *Corvus corone* has been carried out since 1999. The site features a low-intensity agricultural landscape with a mosaic of woodlands, pines and poplars plantations, agricultural fields, uncultivated land, and scattered villages. Here, and unlike elsewhere in Europe, crows exhibit a complex cooperative breeding system where more than two individuals participate in nestling care (Baglione, Marcos, and Canestrari, 2002). Their social structure is based on stable kin groups, typically comprising a dominant breeding pair and non-breeding offspring (mainly males) from previous broods that remain on the natal territory for up to four years (Baglione, Marcos, and Canestrari, 2002). Groups can also comprise immigrants (mainly males) that are related to the territorial breeder of the same sex, and that can share in reproduction (Baglione, Canestrari, Marcos, Griesser, et al., 2002; Baglione, Canestrari, Marcos, and Ekman, 2003). Within groups, relationships are governed by linear and stable hierarchies, where dominance ranks are sex- and age-structured, with breeding males occupying the highest rank, followed by male immigrants, non-breeding male offspring, breeding females, female offspring and lastly, female immigrants (Chiarati et al., 2010). Interactions such as threat postures, allopreening, and food sharing reinforce these social ranks, while conflicts are typically low in intensity (Chiarati et al., 2010). Group members can contribute to chick provisioning, nest cleaning, territory defense, and predator vigilance (Baglione, Marcos, and Canestrari, 2002). Previous studies reveal a finely tuned division of labour and alternation system in chick provisioning within cooperative family groups (Canestrari, Marcos, and Baglione, 2005; Baglione, Canestrari, Marcos, and Ekman, 2006; Baglione, Canestrari, Chiarati, et al., 2010). Breeders and helpers share parental duties, but roles differ subtly according to social category and sex. Breeders contribute more than non-breeders to chick provisioning, and breeding females, which are the only group members that have been observed incubating eggs and brooding chicks, also take on the great majority of nest-cleaning activities (Canestrari, Marcos, and Baglione, 2005; Bolopo et al., 2015). Crows exhibit turn-taking in feeding visits, minimizing temporal overlap between provisioning individuals and minimizing repeated visits (Trapote et al., 2024). Nestling body mass increased with the degree of adults' alternation, likely because well-coordinated groups delivered food at more regular intervals (Trapote et al., 2024). Crow nests can be parasitised by great-spotted cuckoos *Clamator glandarius*, whose chicks are raised alongside crow nest-mates (Canestrari, Marcos, and Baglione, 2009; Canestrari, Bolopo, et al., 2014).

#### *2 Data collection*

All procedures followed ASAB/ABS guidelines (Committee et al., 2024) and Spanish regulations for animal behavioral research, and were approved by Junta de Castilla y León (reference of first released license:EP/LE/177-1999; last released license: EP/LE/681-2019).

The full details on the methodology for determining individual identity is described in Trapote et al., 2024. Briefly, every year all fledglings were captured in the nest and marked with wing tags (Caffrey, 2000). Adult group members were captured with walk-in traps and snap traps, and marked with colored bands and wing tags. In some nests, not all adults were individually identifiable (i.e., 2 or more birds without markers). We collected 50-200 microliters of blood from the alar vein of all banded individuals for molecular sexing and parentage analyses (Baglione, Marcos, Canestrari, and Ekman, 2002). Parentage and sex were determined (Materials and Methods, Section 3). Tables S1-S2 provide a summary of the groups and adult individuals included in the dataset.

We used animal-borne tags to record vocalizations, measure body acceleration, and infer motion behaviors from freely-behaving crows. The full details of hardware and data collection are described in (Baglione, Canestrari, Cusimano, et al., 2025; Stidsholt et al., 2019; Rutz and Troscianko, 2013). The onboard sensors recorded audio, tri-axial accelerometer, and tri-axial magnetometer. We use the term audio-logger for this multi-sensor device. The audio-logger was attached to the two central tail feathers with the stem of a balloon, which eventually degraded, allowing the audio-logger to detach autonomously. Audio-loggers also contained a micro radio transmitter, and were retrieved using a Biotrack Sika receiver. Tables S1-S2 gives a summary of recording durations. Prior work found that the biologgers had negligible effects on brood feeding rates and reproductive success (Baglione, Canestrari, Cusimano, et al., 2025).

We recorded nest camera videos to collect complete information on visits to the nest, including the visits of untagged birds. In 23 out of 25 nests, we filmed nest visit behavior using small camouflaged video cameras from a distance of ca. 1.5–2 m (Canestrari, Marcos, and Baglione, 2005). The nest camera video data did not always overlap with the audio-logger data, nor did different audio-logger deployments always overlap. All sensors recorded time in seconds within a file (as there were typically multiple files per camera or audio-logger) and the sensor time in UTC (local time: UTC+2) but were subject to desynchronization (see e.g., Smeele et al., 2024; Wild et al., 2024). We describe the process for synchronization below (Materials and Methods, Section 8). We defined the time of events and performed synchronization using UTC timestamps.

To evaluate potential sampling biases, we considered our procedures in light of the STRANGE framework for animal behavior research (Webster and Rutz, 2020). Each breeding season, we attempted to locate as many carrion crow breeding groups as possible within our long-term study area (Baglione, Marcos, and Canestrari, 2002) and monitored all nesting attempts, including replacement clutches after early failures. At the start of the breeding season, groups for nest recording and logger deployment were chosen so that the overall sample represented the range of group sizes present in the study population. Nests located in trees unsuitable for camera installation were excluded. Inevitably, some deviations from the intended sampling design occurred due to nest failures or unsuccessful trapping of adults for tagging. Where possible, such cases were compensated by including another group of the same or a similar size (±1 individual). The final sample of tagged birds included in the analyses had a relatively balanced sex ratio (23 females and 20 males), whereas the proportion of breeders and helpers was skewed towards breeders (0.79 vs. 0.21, respectively), mirroring the structure of the study population (Baglione, Marcos, and Canestrari, 2002). Taken together, this evaluation suggests that scope for sampling biases was limited and that the nests included in the final dataset captured the diversity of social contexts present in the study population. Consequently, we expect our results to be broadly representative of cooperatively breeding carrion crow populations.

#### *3 Genetic assignment of reproductive roles and sexing*

Genomic DNA was extracted from blood samples from all individuals sampled using a commercial kit (High Pure PCR Template Preparation Kit, Roche) following the manufacturer's instructions. All individuals were sexed by determining the size of two different introns of the CHD gene. For this, the following primer pairs were used: 3007/3112 (Ellegren and Fridolfsson, 1997) and sex1'/sexmix (N. Wang et al., 2010). PCR conditions available under request.

We screened all samples with the same set of 16 independently segregating polymorphic microsatellites and laboratory procedures used by Trapote et al., 2024 for parentage assignment in the same population. PCR products (1.2 $\mu l$) were mixed with 16 $\mu l$ formamide containing GENESCAN-500 (ROX) Size Standard (Applied Biosystems, ABI) and the allele size of PCR products was determined on a 96-capillary 3730xl DNA Analyzer (ABI). Allele size was independently scored by two researchers using GENEIOUS 8.1.9 (www.geneious.com).

Parentage assignment requires a reference population to obtain allele frequencies. For this, we used the multilocus genotypes of 103 individuals from the same population produced by Trapote et al., 2024. Allele and genotype reproducibility was ensured by (i) using the same genetic analyzer, (ii) including positive controls for each locus in each run, and (iii) repeating the genotyping of 10% of samples for all loci.

Multilocus genotypes were firstly inspected for exact matches. Purging of duplicates left a final dataset of 19 offspring, 52 candidate females, and 60 candidate fathers.

Number of alleles observed, observed and expected heterozygosities, as well as the heterozygosity-based test for Hardy-Weinberg proportions (10,000 permutations) were calculated with GENODIVE 3.03 (Meirmans, 2020). Deviations from Hardy-Weinberg equilibrium were also tested using the probability test (default parameters) implemented in the web version of GENEPOP 4.7 (Rousset, 2008). We adjusted alpha = 0.05 by applying the Holm-Bonferroni correction.

We used COLONY 2.0.6.6 (Jones and J. Wang, 2010), which implements a full-pedigree likelihood approach to simultaneously infer sibship and parentage among a group of individuals provided that multilocus genotype data are available. We performed ten replicate runs for all samples, each of them with a different random number seed. All replicates were run using the following setting: dioecious and diploid species, female and male polygamy, outbreeding model, without clones, medium run length, full-likelihood estimation, medium length of run, allele frequencies estimated from the dataset (no updating), no sibship size scaling, and no sibship prior. Dropout rate, error rate per locus, and the probability of the genotype of a real parent to be present in the multilocus datafiles of candidate adults were set as in Trapote et al., 2024. Number of known paternal sibships, known maternal sibships, offspring with excluded fathers, offspring with excluded mothers, excluded paternal sibships and excluded maternal sibships were all set to zero. Only those cases of parent pair assignment that appeared in at least 7 of the 10 replicates with probabilities $\geq 0.95$ were considered.

We double-checked the parentage assignment for each offspring as implemented in CERVUS 3.0.7 (Kalinowski, Taper, and Marshall, 2007). Simulation parameters were set to: offspring = 10,000, candidate mothers = 52, candidate fathers = 60, proportion of candidate mothers and fathers sampled (0.816 and 0.614, respectively, from Trapote et al., 2024), proportion of loci typed = 0.992, proportion of loci mistyped and error rate in likelihood calculations = 0.035 (from Trapote et al., 2024), minimum number of typed loci = 8). We also used CERVUS 3.0.7 in order to calculate the mean polymorphic information content (PIC) of our set of 16 markers, and the combined non-exclusion probability (parent pair).

All individuals were unambiguously sexed, as both introns produced the same banding pattern for all individuals.

The percentage of missing data in the microsatellite dataset was very low (mean proportion of loci typed = 0.9924). These 16 microsatellite markers showed moderate polymorphism (average number of alleles per locus 6.313, PIC = 0.562), while producing a high degree of confidence in the parentage assignment (combined non-exclusion probability for parent pair = 0.00000013). These values are in line with Trapote et al., 2024, whose simulations showed that this panel of 16 microsatellites produced a high probability (98.1% ± 0.2) of correctly leaving unassigned parentage when the parent has not been sampled; whereas the probability to correctly assign parentage when the parent has been sampled is 97.3% ± 1.9.

Re-screening the electropherograms produced by Trapote et al., 2024 with the new ones generated for the present work, suggested that for marker L14 a pentanucleotide compound repeat may be more accurate than the previously assigned tetrameric motif. Indeed, Fleischer et al., 2008 reported that locus L14 (MJG1; Li, Huang, and Brown, 1997), originally developed for Aphelocoma ultramarina, exhibits a pentameric (GAAAA) motif in the common raven (*Corvus corax*), a species more closely related to *C. corone*, making microsatellite transferability more plausible.

Loci L2, L13, and L19 might deviate from Hardy-Weinberg expectations after correction for multiple testing (L2, probability test, p = 0.0004; L13, probability test, p = 0.000; L19, heterozygosity deficiency, p = 0.0002). Parentage assignment was therefore calculated with COLONY for the whole dataset (16 loci) and excluding these three markers. Results showed no difference in parentage assignment, but probabilities were higher when using the complete set of markers. Additional checks included decreasing the probability of having sampled the biological father and mother (0,5 each) and changing the length of run to “very long”: neither of them modified the parentage assignment nor decreased the reported probabilities.

### *4 Vocalizations*

To chart the repertoire and map its associations to behavior, we required the time of occurrence of vocalizations, their caller, and a representation of their acoustic structure. To represent acoustic structure, we used discrete categories (*call types*) to facilitate the analysis of the entirety of a large corpus of vocalizations. Since we observed that crows often repeat a similar vocalization in quick succession, we grouped vocalization events into *bouts* to improve statistical independence of our dataset.

Crow vocalizations occurred sparsely within multi-hour audio recordings, along with self-movement noise (e.g., wings flapping, splashing), environmental sounds (e.g., wind) and the vocalizations of other species. To obtain vocalization events from this continuously-recorded audio, we first used supervised machine learning to detect the vocalizations in time and make an initial caller assignment (as *focal*, i.e. made by the adult bird wearing the audio-logger; or *non-focal* adult or chick) (Baglione, Canestrari, Cusimano, et al., 2025). Then, to develop a set of call types, we used a machine-learning based clustering of denoised focal vocalizations to guide manual annotation of call types (see similar approaches in Thomas et al., 2022; Merino Recalde, 2023). Using supervised machine learning trained on these annotations, we assigned a call type to every focal vocalization. After synchronization (Methods and Materials, Section 8), we post-processed the vocalizations to refine the caller based on multiple aligned audio channels, when possible. Finally, we assigned each vocalization to a bout based on the time between vocalizations of the same type. Focal vocalizations by individual and call type are summarized in Tables S1-S3 and Figure 2B. For all vocalization processing except denoising (which required a sampling rate of 22050 Hz), we down-sampled the audio at 16 kHz.

#### Detection and initial caller assignment

We detected vocalizations as described in Baglione, Canestrari, Cusimano, et al., 2025, which we summarize here. One of the authors (DC) used Raven Pro (K. Lisa Yang Center for Conservation Bioacoustics at the Cornell Lab of Ornithology) to annotate 35 audio files from 14 birds tagged in 2019, with a total duration of 187.7 hours. Vocalizations were annotated with a bounding box indicating start- and stop-time, as well as caller identity. Bounding boxes could overlap in time. For caller identity, we used four categories: focal adult, non-focal adult, crow chick, or great-spotted cuckoo chick. Vocalizations could also be annotated as Unknown, if it was difficult to determine whether the caller was focal or non-focal. We then trained an early version of *Voxaboxen* (Mahon et al., 2025), an onset-based sound event detection model, to predict start-time, stop-time and caller identity. We filtered out detections with predicted detection probability less than 0.5, and marked all detections with class probability less than 0.5 as "Unknown". On held-out individuals, this model achieved a precision score of 0.78, a recall score of 0.77, and F1 score of 0.78 (scores averaged across the four target categories). The model obtained slightly better metrics for focal vocalizations (precision: 0.83, recall: 0.79, F1: 0.81). We applied the trained model to all recorded audio, and obtained a list of vocalizations across all audio files. This resulted in 127750 detected focal vocalizations.

#### Denoising

We aimed to use automated clustering on the focal vocalizations to guide manual call type annotation (see below), but found that background audio had a strong influence on clustering. Therefore, we performed a denoising procedure to remove these sounds from the focal vocalizations, using the *BirdMixIT* source separation model (Denton, Wisdom, and Hershey, 2022). BirdMixIT is a time-domain convolutional neural network that takes a single-channel signal containing a mixture of sounds as input, and predicts the set of pre-mixture sounds which created this signal (*stems*). BirdMixIT is trained on mixtures of soundscapes which include bird vocalizations, and we applied it to the carrion crow focal vocalizations without fine-tuning. Since BirdMixIT outputs unlabeled stems (i.e., it does not indicate which stem contains the focal vocalization), we used our trained detection model to select the stems containing the denoised focal vocalization.

Specifically, for each focal vocalization, we took a clip beginning 2.5 seconds before the focal vocalization, and ending 2.5 seconds after the focal vocalization. We applied the pre-trained BirdMixit source separation model to this clip, which resulted in four stems. To identify the stem with the focal vocalization as the main sound, we used our trained detection model. For each stem, we computed a confidence score $C$ based on the detection model's outputs: $C = \max D_i(p_v(D_i)/p_f(D_i))$, where $D_i$ are the focal vocalizations detected beginning between 2-3 seconds, $p_v$ is the Voxaboxen probability that the detection is a vocalization, and $p_f$ is the Voxaboxen probability that the vocalization is from the focal bird.

We then took the stem with the highest confidence score, to use in subsequent steps. We tested this stem selection procedure using 90 vocalizations (15 clips each from 6 birds). One of the authors (MC) annotated the stem that appeared to have the most sound energy coming from the focal vocalization. The procedure correctly selected the annotated stem 95.5% of the time. A small percent ($< 5\%$) of the time, the detection model predicted that there were no focal vocalizations in any of the stems. In these cases, we used the original, noisy sounds in subsequent steps to ensure that we did not systematically miss any vocalization types.

#### Call type annotation

To determine the call types, we guided manual annotation using initial clusters that were based on a self-supervised machine learning representation. We first obtained a subset of the sounds that would be clustered and then considered for manual annotation: we subsampled 9000 focal vocalizations from the 127750 originally selected (7%). To do so, we embedded the set of denoised focal vocalizations with AVES, a transformer-based audio representation model (Hagiwara, 2023). This model converted each vocalization into a 768-dimensional vector. We clustered the embedding vectors using k-means clustering (k=300), and then we sampled 30 vocalizations uniformly at random from each cluster. Clustering before sampling was performed to obtain examples of rare but acoustically different vocalizations, which would likely not be selected if we sampled uniformly from the entire dataset. Qualitatively, we found that these clusters often over-separated similar common call types (like caws and grunts, described below) and grouped together acoustically distinct rare call types. We ultimately annotated 7443 vocalizations due to time limitations; a review of the remaining 1557 did not surface any new call types.

After an initial review of the data, we attempted to increase the size of initially assigned call types with fewer than 20 examples, in order to enable accurate classification on the rest of the dataset. We computed audio features using a pre-trained BEATs encoder (Chen et al., 2022) for all sounds in the dataset, padding each sound with zeros to a duration of 2 seconds and averaging the embedding vectors over time. For each initially assigned call type with less than twenty examples, we then obtained its 50 nearest neighbors by retrieving unannotated vocalizations with the minimum L2 distance to any annotated example. One of the authors (CG) used a custom interface to manually select any of the nearest neighbors that matched the initially assigned call type. In total, this process added 420 annotated vocalizations.

The final total number of annotated vocalizations was 7863. The remainder were assigned to call types by the call type classification model (see next section).

One of the authors (CG) manually reviewed the subsampled calls using custom software that represents the calls in a 2D acoustic space and allows one to assign each call to different groups. This resulted in the initially assigned call types mentioned above. A second author (BH) iterated on this method, further combining similar groups. Other authors (MC, VB, DC) verified the groupings throughout the process. Vocalizations were grouped based on listening to recordings, as well as visually inspecting the spectrograms, with attention to overall similarities in duration, pitch, pitch contour, noisiness/harmonicity, amplitude, amplitude modulation/trilling, and presence of clicks (presumably made by bill-clacking). This resulted in 3 super-categories of vocalizations: Caws, Grunts and Exceptional. The majority of calls were assigned to one of two super-categories (Caws and Grunts), which contained 2037 and 2137 vocalizations, respectively. In both of these, there was considerable variation in the acoustic features of the vocalizations, but the variation was continuous and there was no clear way to further subdivide the families. The Exceptional super-category consisted of 3390 vocalizations that had been manually organized into 22 call types that were characterized by distinctive acoustic features or structural elements; each of these is identified with a randomly assigned call type number (i.e. a numerical hash). Text descriptions of caws, grunts, and all exceptional call types are given in Table S4. Initially, 11 additional call types were formed from 51 vocalizations, but each contained fewer than 20 vocalizations and were discarded from the call type classification scheme (these vocalizations were each later re-assigned to one of the other call types by the call type classification model; they should add minimal noise to analyses because they represent $< .04\%$ of the overall dataset). Finally, 248 vocalizations were annotated as non-adult crow vocalizations or ambient noise (false-positive detections).

#### Call type assignment

We trained a machine learning classifier to classify vocalizations into Caws, Grunts, or an Exceptional call type. This was a 25-way classification problem (2 super-categories, 22 exceptional call types, 1 non-adult crow). The annotated calls were randomly divided into Train/Val/Test splits (ratio 0.7/0.1/0.2), stratified across call types. We used macro-averaged F1 score to measure performance.

Each vocalization was right-padded and cropped to a total duration of 1 second. The classifier used the pre-trained BEATs base model (Chen et al., 2022) to extract features (768 dims, 50 Hz frame rate) from this audio. These features were mean-pooled in the time direction and passed through a MLP (1 hidden layer, 1024 dims).

The model was trained using categorical cross entropy loss for 100 epochs, using a cosine learning rate scheduler, batch size 16, and the Adam optimizer (Kingma and Ba, 2014). Throughout training, the BEATs feature extractor was frozen. We swept learning rates from 3e-5, 1e-4, 3e-4 and selected the model with the best macro-averaged F1 score on the Val set. (To ensure that the model made predictions based on the characteristics of the vocalization, and not the background noise, we sampled audio clips from from intervals of audio-logger data that did not contain detected adult crow vocalizations. For each training example, we summed the waveform of the crow vocalization with the waveform of the background audio.)

On the Test set, the classifier obtained a macro-averaged F1 score of 0.924. Per-call type, the classifier's performance ranged from F1 of 0.658 for the non-adult crow call type to F1 of 1.0 for four different call types. The low performance on the non-adult crow call type is likely due to crow chick begging vocalizations that the classifier confused with adult crow vocalizations, since these can be acoustically similar. For all other call types, F1 was at least 0.800.

We applied the call type classification model to all of the previously detected focal vocalizations. There were 34768 caws, 53685 grunts, and 28851 vocalizations from the exceptional call types (125188 total). We omitted from further analysis all 2508 vocalizations classified as non-crow. To probe whether calls with continuous acoustic variation vary in terms of their associations with behavior, we split the graded super-categories into call types based on amplitude and duration quantiles. For each of the two graded super-categories (caws and grunts), we first subdivided it into six call types based on the duration quantiles (0.2, 0.4, 0.6, 0.8, 0.9, 1.0) Figure S8 denoted as Caw0, Caw1, Caw2, Caw3, Caw4, and Caw5 and similarly for Grunt; the 0.9 quantile was added because the distribution of call durations was right-skewed. We chose duration as a primary method of division because it was easy to measure and its variation was apparent in visual inspection of spectrograms. Since the number of grunts was approximately 1.5 times the number of caws, we additionally subdivided each duration quantile based on the median amplitude value over all grunts, resulting in twelve categories (Grunt[0-5]Q denoting the lower-amplitude half, and Grunt[0-5]L denoting the higher-amplitude half), with amplitude being measured from the denoised vocalizations where possible. We chose amplitude because it is easy to measure, its variation was apparent in visual inspection of spectrograms, and we reasoned that continuous variation in amplitude may be more relevant for grunts because they have a relatively low mean amplitude level. In the final focal vocalization dataset, the most frequently-occurring exceptional call type (E4) occurred in 18369 calls (16.0%) and the least frequent (E47) in 119 calls (0.1%) (Figure 2B; see Table S3 for a full list).

#### Caller identity annotation with synchronized audio

Our initial caller assignments were based on annotations on audio from a single audio-logger. Post-synchronization, we were able to compare detections registered by multiple audio-loggers to refine our caller annotation. We followed a similar procedure to Zeh et al., 2024, comparing across multiple audio-logger channels to uniquely assign a focal vocalization to a caller. The first step was to match any detections across channels stemming from the same vocalization, and then to select which bird was focal (if any) via a relative amplitude comparison. The size of our dataset required that we automate the first step, so we annotated a dataset to enable us to match vocalizations across audio-loggers.

From each nest with multiple synchronized audio-loggers, we sampled synchronized audio clips from multiple individuals (details on sampling below) and used Audacity (Audacity Team; version 3.7 or 3.8) to view and annotate multiple clips simultaneously. As in the single-channel protocol described above, we annotated clips with bounding boxes corresponding to the start- and stop-time of vocalizations, and with caller assignments. We retained the caller categories of focal, non-focal, crow chicks, cuckoo chicks, and Unknown. In addition, we either annotated each bounding box as *alone* indicating it was only registered in a single channel, or provided it with a (non-unique) event label that had a *match* in at least one other channel, to indicate that it was the same vocalization registered in multiple channels. When available, we watched the corresponding nest camera video in case it provided additional cues to caller identity. The single-channel detections were also presented in Audacity, as a visual guide to speed annotation, but annotators were not constrained to use them. Annotations underwent automated review for internal consistency, such as only having one focal caller across matched detections.

We annotated in two sets, using the first to evaluate interrater agreement. To increase efficiency, we used the existing single-channel detections to select clips likely to contain matched vocalizations. We found all 1-minute intervals where focal detections from different audio-loggers occurred within 1.5 sec of each other, discarded intervals not meeting this criterion, and combined consecutive intervals into clips. In the first set, we selected 2 to 4 clips for each nest available. This resulted in 18 clips across 8 nests (37 min total, min clip duration=1 min, max clip duration=6 min, mean number of focal detections per clip=79.5). In the second set, we sampled for additional variety. For each nest, we selected the 5 clips with the highest rate of focal detections, sampled 2-3 clips from the 10 with the highest rate of chick vocalizations (to obtain clips likely to be at the nest), and did the same for non-focal vocalizations. This resulted in 109 clips across 12 nests (184 min total, min clip duration=1 min, max clip duration=11 min, mean number of focal detections per clip=54.7), including at least three minutes of audio for each available individual and at least five minutes of audio for each available nest.

Two annotators (authors AM and MC) annotated the first set. To assess interrater agreement, bounding boxes were matched based on their intersection-over-union (IoU threshold=0.5, with bipartite graph matching (Mahon et al., 2025)). There were 2604 events annotated by both authors and 468 vocalizations annotated by only one (84.8% agreement). Of the former, annotators provided the same caller label 87.0% of the time and the same event label 87.0% of the time. Then, author AM annotated the second set. For the second set, any chick annotations less than 5 seconds apart were combined to reduce annotation time. In training and evaluation of our machine learning model (next section), we used both sets (the two sets were non-overlapping except for one clip, which was discarded from the first set).

#### Caller identity assignment with synchronized audio

To automate matching vocalizations across channels, we trained a machine learning model to classify a pair of vocalizations as a *match* or not. We then used this model to greedily assign detections across channels to shared vocalizations events (or label them as *alone*). Last, we decided which of the detections were focal using relative amplitude.

The annotated files were randomly divided into Train/Val/Test splits (ratio 0.7/0.1/0.2), stratified across nests. All pairs from detections within 1-sec of each other were created, and the annotations indicated whether they were matched or not. 3-sec clips centered on each detections were generated and the clips of a pair were concatenated. On each training epoch, a subset of all pairs was randomly sampled, subject to balancing the nest and matched vs. non-matched pairs.

The classifier used the pre-trained BEATs base model (Chen et al., 2022) to extract features (768 dims, 50 Hz frame rate) from this concatenated audio. The intervals of the embedding corresponding to the detections were extracted. Each extracted interval was mean-pooled and then these average embeddings were concatenated along with the durations of the original detections (1538-dimensional feature vector). A linear classifier predicted logits for whether the pair was matched or not.

To reflect our use case at inference time (see below), the evaluation metric on the Val and Test sets was the F1-score of the focal caller assignments, computed across detections in a focal/non-focal match. To obtain this metric, we post-processed the logits to create matches greedily. If a pair of vocalizations had a logit above threshold (determined by hyperparameter search), they were paired, subject to the constraint that matches do not contain more than one detection from one channel. If a detection was not assigned to any match, it was considered alone. Then, for all matches, we assigned the caller as the focal individual with the highest root-mean-square amplitude from the onset to offset of the detection. If the model correctly assigned that detection as alone, then this would not affect the evaluation metric (because only matched detections are used to compute the evaluation metric). However, if the model incorrectly assigned this detection to a match, then it would decrease the F1-score.

The model was trained using binary cross entropy loss for 50 epochs, batch size 5, and the Adam optimizer (Kingma and Ba, 2014). The BEATs feature extractor was unfrozen on epoch 3. We swept learning rates from 1e-5, 3e-5, 5e-5, logit thresholds from -4.0, -3.0, -2.0, -1.0, 0.0, 1.0. We selected the model with the best focal F1 score on the Val set (learning rate = 5e-5, logit threshold = -4.0).

On the Test set, we obtained a focal F1 score of 0.91. Per nest, the performance ranged from F1 of 0.61 for nest 2019-EB, to F1 of 1.0 for nests 2018-AW, 2019-O, and 2019-CV. To understand baseline performance, we also evaluated an oracle model, which used the ground-truth matches and the relative amplitude heuristic to select a caller: its F1 score was 0.93.

To apply this network to the original detections, we found vocalizations which occurred within 2-sec of vocalizations in another channel, and paired them. We generated matches as described above. If a detection was not matched, it was considered *alone* and retained its original caller assignment. If detections were matched, then their original caller assignments were considered. Of the original assignments, (1) if there was at least one crow or cuckoo chick, all changed to that chick label, (2) if all were non-focal, they were not changed, (3) if there was only one focal, they were not changed, (4) if there was more than one focal, a new unique focal vocalization was assigned according to relative amplitude. This resulted in 11058 originally focal vocalizations that were re-assigned to crow chicks, cuckoo, or non-focal. The final number of focal vocalizations was 114676.

#### Bout assignment

We observed that crows made sequences of repeated calls. For each call type, we first plotted the kernel density estimate of the log10-scaled time between two calls of the same type, by the same caller (Figure S9). We found a distinctive distribution shape across multiple types, which had more than one mode: a strong first peak typically around 1 sec, followed by a local minimum and weaker peak. The major exception to this pattern was for the grunts super-category, and some call types had a weak second peak. Given the qualitative similarity of the distributions, for each, we estimated the location of the local minimum of the kernel density estimate (using functions stats.gaussian_kde and signal.find_peaks with the default settings in scipy v1.16.0). The median of the local minimums across call types was 5.12 sec, which we set as the bout threshold. We then assigned vocalizations (with the same call type and caller) to the same bout when the time between them was less than the bout threshold. The number of vocalizations per bout for each call type is plotted in Figure S10.

#### Call type amplitude

We provide relative measurements of vocalization amplitude since reference to an absolute external sound pressure level was unavailable, but microphone sensitivity was calibrated to within ±3 dB across sensors. For each vocalization, we measured its root-mean-square amplitude (using denoised vocalizations where possible). We then computed decibels relative to full scale (i.e., reference = 1) for each sound. We took the mean and standard deviation within a call type category (Figure 2B, black dots). We found the range of the call type amplitude means was approximately 30 dB, much greater than between-sensor variation. We also computed the relative amplitudes of audio clips containing crow chick vocalizations while adults were visiting the nest (n = 668, across 34 audio-loggers; Figure 2B, horizontal line).

### 5 *Flights, high-activity, and low-activity intervals*

From the audio, we identified intervals when birds were flying or not flying (Stowell, Benetos, and Gill, 2017). After nest visit annotation, we additionally defined flights when birds arrived at or left the nest, since we hypothesized they may be particularly associated with coordination-relevant vocalizations. We split the remaining flights into short and long flights. After synchronization of audio and nest visits, we identified intervals when the bird was neither flying nor at the nest. We used the accelerometer data to subdivide these intervals when the bird was neither flying nor at the nest into *high-activity* and *low-activity* intervals. In summary, we used the logger data to define arrival flights (n = 3060), departure flights (n=3050), short flights (n = 9063), long flights (n = 7887), low-activity intervals (n = 111259), and high-activity intervals (n = 134568).

#### Flight annotation

We sought to automatically detect flying behavior from the audio-logger data. We defined a binary ethogram (*flying* or *not flying*), with frames left unannotated being considered as *unknown*. *Flying* was associated with the sound of wing flapping and wind, and often contained a high-amplitude, periodic (approx. 5 Hz) signal in the accelerometer channels (these periods could contain flying without wing-flapping, i.e., gliding/soaring). *Not flying* included a variety of behaviors such as wing flapping without moving, hopping, preening, bathing, walking, resting, and chick feeding.

Authors MB, AM, and MC used Audacity to annotate an initial set of 32 2-minute clips of accelerometer and audio data; qualitative discrepancy was only found in one file due to a potential gliding interval. We then sampled a final set of clips for annotation. We sampled five two-minute clips from each tagged individual in the dataset, for a total of 165 clips. These clips came from 145 unique audio-logger files. We designed the sampling procedure such that for 12% of clips, the clip contained at least one nest visit. Author AM annotated 109 clips and author MB annotated 87 clips, with 31 clips being the same for the two annotators so that we could evaluate interrater agreement. On the shared 31 clips, a small proportion of the total data labeled by either annotator as *Not flying* ($< 5\%$), was labeled by one of the annotators as *Unknown*. Disagreement on *Flying* vs. *Not flying* amounted to less than 10 seconds.

#### Flight detection

Using the annotated data, we trained a flying detection model. We partitioned the data into Train/Val/Test sets, splitting on the level of individuals, at a ratio of 0.6/0.2/0.2. We measured performance using the frame-wise F1 score. For the purpose of evaluation, predictions for the frames annotated as Unknown were ignored.

The model used a BEATs (Chen et al., 2022) encoder, followed by a linear prediction head with a sigmoid activation function, with two outputs. Predictions were made at a 50 Hz frame rate. The two outputs indicate the probability that the bird is Flying in a given frame, and that the bird is Not Flying in a given frame. When the model predicted that the probability of Flying was >= 0.5 for a given frame, we interpreted the output as Flying = True, and similarly for Not Flying. At inference time, when the model predicted Flying = Not Flying = False for a frame, or when the model predicted Flying = Not Flying = True for a frame, we marked that frame as Unknown; however, in practice we found that this second situation did not occur.

The model was trained with binary cross entropy loss for 50 epochs, with batch size 8, with the Adam optimizer (Kingma and Ba, 2014) and a cosine learning rate scheduler. Loss was masked for frames annotated as *Unknown*. During training, the pre-trained BEATs encoder was frozen for the first three epochs and then unfrozen for the rest of training. As an initial step, we trained models using learning rates selected from (1e-5, 3e-5, 1e-4) on the Train set. The learning rate 1e-5 led to the model with the best average frame-wise F1 score on the Val set. Then, we trained a model using the combination of the Train and Val set, using this learning rate. The final model obtained an average frame-wise F1 score of 0.918 (Flying = 0.907, Not Flying = 0.930) on the Test set.

We applied the trained model to the entire dataset. In doing so, we applied some post-processing steps. First, we smoothed the binary predictions using a max-pooling filter, followed by a min-pooling filter (diameter 25 frames for each). Therefore intervals with $< .5$ second gap were merged, and remaining intervals $< .5$ seconds were removed. Finally, we manually inspected one file to confirm that the predicted Flying intervals were accurate.

#### Flight types

After synchronization of the nest visits and flights (Materials and Methods, Section 8), we found flights that directly preceded or followed the nest visits. We defined *arrival flights* as the flight immediately preceding the nest visit start. We defined *departure flights* as the first flight with its start after the nest visit start and its end after the nest visit end. Birds typically interacted with chicks soon after landing at the nest, meaning that typically the end of arriving flight and the beginning of nest visit occurred in close succession. Sometimes birds remained near the nest after interacting with chicks but were not on camera (one criterion for visit annotation, see Materials and Methods, Section 7), such that leaving flights did not immediately occur after the end of a visit.

Flights that were synchronized with nest visits, but were not an arrival or departure flight, were labeled with destination *Other*. We split *other* flights into two categories, *short* and *long*, based on the median duration of all flights (10.02 sec). Flights that were not synchronized to nest visit data were not included in Analyses 2-4 (Materials and Methods, sections 11-12), which required specific flight types.

#### Low- and high-activity intervals

To obtain low- and high-activity intervals based on the accelerometer data, we measured the overall dynamic body acceleration (Wilson, White, et al., 2006) and then characterized not-flight/not-visit intervals as low or high-activity based on a threshold. We used overall dynamic body acceleration because it is correlated with other metrics of overall activity level (Wilson, Börger, et al., 2020; Elliott, 2025) and an effective feature in coarse-grained behavior classification (Nathan et al., 2012), including in carrion crows (Hoffman et al., 2024). We intended to make a coarse distinction between high and low activity rather than obtain precisely calibrated estimates of energy expenditure.

First, we normalized the average field strength within each raw accelerometer file to be equal to 1 g. Then, for each raw tri-axial acceleration channel, we applied a high-pass delay-free filter. We used a linear-phase symmetric FIR filter with a Hamming window, followed by group delay correction (Ruiter et al., 2020). The high-frequency component is termed the dynamic acceleration, as filtering attenuates the acceleration due to gravity. The filter cutoff was set to 3 Hz, based on visual inspection of the accelerometer time series (Baglione, Canestrari, Cusimano, et al., 2025). Then, in non-overlapping one-second intervals, we measured the mean overall dynamic body acceleration (ODBA) – the sum of the absolute value of each channel of dynamic acceleration. We found the median value of the mean ODBA in non-flight intervals that occurred during the daytime (median = 0.094 g; see Figure S11), and set this as our threshold. Then, we categorized each one-second interval as below or above threshold. To bridge short changes in category to give more contiguous intervals, we used a median filter (width = 5 seconds) on the binary time-series of categorized one-second intervals (Cakir et al., 2015). Finally, we combined any contiguous intervals with the same categorization to produce the final low- and high-activity intervals.

### *6 Video annotations*

We manually annotated videos and audio-logger data to determine nest visits and to synchronize the sensors. We first provide an overview of both of these annotation types since they were done simultaneously with video, then give full details on each separately in sections below. Nest visits were annotated as a time interval when a particular individual arrived at and then departed from the nest. To synchronize sensors, we annotated synchronization *keypoints*: timepoints of events detectable in both video and audio that could be used to align the files. We annotated videos that overlapped with audio-logger data from at least one bird. Annotators also noted other significant events: predation, other chick deaths, fledging, and events affecting data quality such as premature audio-logger detachment.

First, videos from 2019 were manually annotated with nest visits and keypoints. Each nest was fully annotated by one of the two annotators (author AM or MB). Then, we performed an interrater agreement analysis on a subset of videos from 2019. This revealed sufficiently consistent nest visit annotations and synchronization. All the remaining nests (2018 and 2021) were annotated with nest visits and keypoints, each by one of the two annotators.

Annotations were made using Premiere Pro (Adobe; version 24.5.0, Build 57). The Premiere Pro timeline interface could contain multiple channels of data: video (image and audio), logger audio from multiple birds, and a visual indicator for intervals likely to contain nest visits for each tagged bird (derived from chick vocalization detections; see details below). Markers were placed and labeled to indicate nest arrivals, nest departures, and keypoints, while the color of the marker indicated the associated individual. For example, an orange marker labeled "a" would indicate an arrival from the bird with orange wing tags. We exported the markers using Visual Studio Code.

#### *7 Nest visits*

The dataset was collected during breeding season, and thus, visiting the nest for chick provisioning was a key joint activity for the crows at this time. As described in Materials and Methods Section 1, crows show alternated turn-taking at the nest which has fitness benefits (Trapote et al., 2024). To reflect this prior work on turn-taking as a measure of active coordination (Trapote et al., 2024), visits were *repeated* if the previous visit was by the same bird, or *alternated* if not. Table S2 gives a summary of nest visits by individual.

For each nest visit, we recorded the identity of the individual visiting, as well as the time of their arrival and departure. Due to the variation in camera and nest geometry, the cues to a bird's arrival and departure varied. Arrivals were defined as when the (1) bird first entered the video frame, (2) when the bird first touched the nest, or (3) when it first interacted with the chicks. Departures were defined by when (1) the bird left the video frame for the last time that visit, (2) when the bird last touched the nest or (3) when the bird ceased to interact with the chicks. If the bird was never visible during a visit, the visit could be determined based on other visible events as well as video/logger audio.

Identity could be inferred from wing tags or leg bands when visible, or the presence of common events in the video and logger audio. Untagged birds were labeled as 'untagged adult', with a numerical index to distinguish untagged individuals if multiple were present at the nest at the same time. Untagged individuals could not be consistently identified across visits. In one case, the identity of a bird visiting the nest could not be determined so it was labeled as 'unknown adult'.

To aid the annotators in finding nest visits efficiently, we obtained crow chick and cuckoo detections from our detection model for each audio-logger (Materials and Methods, Section 4). We transformed these detections into a binary sequence (1 indicating detection) of duration equal to the logger audio, which could be aligned with the video. These detections provided a visual cue for intervals that were likely to contain nest visits, since chick vocalizations may be recorded if an adult is at the nest. Annotators were still required to scroll through the video to identify and verify nest visits by visual inspection, including in periods without detected chick vocalizations.

Annotations underwent automated review for internal consistency, specifically that a bird's arrivals and departures alternated (as arriving or departing multiple times in a row would not be possible).

##### Interrater agreement

We evaluated interrater agreement after completing the annotation for nests in 2019. We selected a subset of videos from 2019 upon which to evaluate interrater agreement. For each nest in 2019 (except CA, which had a short period of audio-video synchronization), we randomly sampled a time of day. We selected all videos overlapping with or after this time until reaching at least eight nest visits from any bird (according to the original annotator). This resulted in a different number of videos for different nests, depending on the rate of nest visits (min=1, max=3). An interval was kept if: (1) it overlapped with at least one hour of logger audio from at least one bird and (2) the interval was between the first and last annotated nest visit of the day, not counting birds who stayed overnight. The second constraint avoided night time intervals when there is almost no activity. This sampling procedure resulted in 23 hours of video. Each annotator annotated the nests for which they were not the original annotator, and the annotator annotated the full duration of each video.

We automatically matched annotators' nest visits based on their intersection-over-union (IoU threshold=0.5, using the bipartite graph matching method from (Mahon et al., 2025). 109 visits were automatically matched and an additional 25 visits from either annotator were not, for a total of 134 visits. Out of the 109 automatically matched visits, 108 had the same identity (one error due to keystroke mistake). We then visually inspected the remaining visits. We found that 13 of these events had correspondences between annotators, but were not automatically matched due to differences in duration. For the remaining 12 events which had no correspondences between annotators, we found 2 were due to different usage of arrive/depart criteria (for 2018/2021, visibility in the frame was explicitly preferred for ambiguous cases), 8 were associated with challenging situations (brief visits, many birds present at the nest, hardly visible bird) and 2 had no associated ambiguities or challenges.

We also quantified the agreement between the timing of arrivals and departures. For this, we only looked at 109 automatically matched visits according to the IoU criterion, and computed the absolute difference in timing for each event. Across all visits, the median absolute time difference in the arrival was 0.40 seconds and the difference in the departure was 0.20 seconds, with these medians corresponding to less than one percent of the averaged event duration.

### *8 Synchronization*

To obtain synchronized events, we annotated synchronization *keypoints*: timepoints of events detectable in multiple channels that could be used to align the files. We used these timepoints to define a function which could map between the time as indicated by the clock on different sensors. We first annotated keypoints in nest camera video and the logger audio for each nest (briefly mentioned in Materials and Methods, Section 6). Audio-loggers of all tagged birds in a nest were synchronized to the video when available. Some audio files occurred in times when no video was available and when multiple audio-loggers were recording. To synchronize these recordings, we separately found keypoints registered by multiple audio channels. After synchronization, we estimated the typical synchronization error, accounting for both annotation and interpolation error, to be sub-second.

#### Audio-video synchronization keypoint annotation

For audio-video synchronization, authors AM and MB selected events detectable in both the video and in the audio-logger. To include an event as a synchronization keypoint, adult birds had to be partially visible in the video or indicated by another visible event (e.g., branch swaying upon landing). A variety of audible events could be used as keypoints, including crow chick vocalizations, cuckoo chicks vocalizations, adult vocalizations, chick feeding, landing, and taking off. We aimed for at least 3 keypoints per video/audio-logger file pair. We prioritized events that were audible in multiple audio-loggers, when multiple birds were at the nest. The video cameras for nests BQ and FA in 2021 did not have audio, so movements visible in the video were synchronized to the audio. This resulted in 8753 keypoints, covering 442 audio-logger files and 997 video files, with an average time of 5.9 min between keypoints in a file.

We evaluated interrater agreement using the dataset described above for evaluating interrater agreement of nest visits. Since annotators were not constrained to choose the same keypoints, we instead looked at the time agreement between detected vocalization events that were synchronized using each annotator's keypoints. We used the keypoints to define a piecewise linear function (using interp1d from scipy.interpolate, v1.16.0) to map vocalization events from audio-logger time to video time. We computed the absolute difference between the timing of vocalizations as synchronized with the two annotators' keypoints. The mean across vocalization and nests was 0.115 seconds (range=[4e-4, 0.32] sec by nest) and the maximum across vocalizations averaged across nests was 0.35 seconds (range=[0.04, 1.14] sec by nest). Last, there were three nests where more than one bird had vocalization detections and there were over 30 vocalizations for each bird. We applied Spearman's rank correlation to check if the vocalizations occurred in the same order given the two sets of keypoints. We found that the two orderings were highly or perfectly correlated ([0.99, 1.0, 1.0], $p < 0.01$ for each).

#### Audio-audio synchronization keypoints annotation

Some remaining audio-logger files only occurred simultaneously with other audio-logger files (not video). To synchronize the audio-loggers to each other directly, we annotated keypoints across multiple audio files. One annotator (author AM) selected keypoints in this data, using Audacity to display multiple audio files simultaneously. Sounds seeming to arrive at the audio-loggers simultaneously were prioritized. We aimed for two annotations per hour per audio-logger pair. This resulted in 773 keypoints, covering 146 audio-logger files, with an average of 26.8 min between keypoints within a file.

#### Synchronization function

We used the keypoints to define a piecewise-linear function mapping between the times in one sensor and the other; this function allowed us to synchronize events across sensors that were not annotated as keypoints. In detail: when synchronizing an event $e_0$ occurring in file $f_0$ recorded by sensor $s_0$ to time as recorded by sensor $s_1$, we first selected all keypoints in file $f_0$ which also occurred in any file recorded by sensor $s_1$. We then defined a piecewise-linear function where input values corresponded to UTC timestamps in $s_0$, output values corresponded to UTC timestamps in $s_1$, and endpoints of the linear segments were the keypoints. Before the first selected keypoint and after the last selected keypoint, we extrapolated assuming no additional drift (slope=1). If no keypoints in $f_0$ occurred in $s_1$, then we listed the timestamp of $e_0$ in $s_1$ as "unknown". Pairs of events which lacked timestamps in a common sensor were left out of Analyses 2-4 (Materials and Methods, sections 11-12).

The absolute difference between the timestamps of keypoints across two files was on average 358.4 sec, the median across individuals (N=37) of this average was 74.3 sec, the standard deviation across individuals of this average was 689.46 sec, and the max across individuals of this average was 4016.10 sec. Therefore, typically events were initially desynchronized by one or more minutes, but this varied by individual.

#### Validation

We estimated the relative synchronization error for non-keypoint events, by leaving out sets of keypoints and measuring the difference between their interpolated time and their annotated time. For each pair of files with at least four keypoints in common (N=1057 pairs), we formed the piecewise-linear interpolation function based on *only* the second and second-to-last keypoints in the file (i.e. holding out the other keypoints). Then, we measured the absolute value of the difference between the interpolated timestamps, and the actual timestamps, of the held-out keypoints. We averaged these errors within each file. The median error across files was 2.6e-6 sec, the standard deviation was 0.62 sec, the 95th percentile was 0.19 sec, and the maximum was 14.7 sec. Therefore, our synchronization procedure successfully decreased the error for non-keypoint times.

### *9 Repertoire-mapping overview*

We first characterized crow daily vocal activity with respect to coarse-level temporal and social factors. Diel patterns in the usage of calls are relatively easy to obtain, and yet they can strongly suggest function (e.g., territorial vocalizations at the beginning/end of daily activity; Cramp, Perrins, and Brooks, 1994). Furthermore, some call functions may be characteristic of specific social roles (e.g., dominance signals; Tibbetts, Pardo-Sanchez, and Weise, 2022) or group composition (e.g., call exchanges during pair formation and nest building; Gill et al., 2015). In carrion crows specifically, such patterns could emerge as adults coordinate to fulfill provisioning needs that are determined by brood characteristics (Trapote et al., 2024).

We next investigated how behavioral states co-occurred with vocalizations. Signallers may use different vocalizations based on their own ongoing state, as in distress signals when the signaller is under threat (call, given signaller state; Mas and Kölliker, 2008), or based on another individual's state, as in courtship displays modified based on the receptivity of the partner (call, given other state; Kelso and Verrell, 2002). Accordingly, the signallers' calls provide information for others to make inferences about the current state of the signaller (signaller state, given call) or a third individual (other state, given call). Such analyses are mainstays of animal communication research (Bradbury, Vehrencamp, et al., 1998); here, we systematized this classic approach into a matrix of associations for the full repertoire.

Finally, the granularity of the dataset allows for an analysis that goes beyond the co-occurrence of vocalizations and states, to consider the precise order in which different events happen. This in turn suggests a systematic typology of associations, with each corresponding to possible functional interpretations. For example, if an individual vocalizes after a behavioral event, this might be interpreted as providing an update on the signaller's state, such as in contact calling after movement (behavior → self vocalization; Kondo and Watanabe, 2009). In contrast, vocalizing before a behavioral event might be interpreted as an announcement of the signaller's intent to act, such as in offensive threat signals (vocalization → self behavior; Waas, 1991). Such relationships may also involve different individuals, as when an arrival is met with a greeting (behavior → other vocalization; Eleuteri et al., 2024) , mobbing calls provoke anti-predator behavior (vocalization → other behavior; Magrath et al., 2015), or de-escalation signals deter aggressive interactions (vocalization → other behavior; Waas, 1991). When both events are vocalizations, these relationships may suggest order in vocal sequences (vocalization → self vocalization) or conversations, as in duetting (vocalization → other vocalization; Hall and Magrath, 2007). The resulting matrix of vocalization and behavior pairs, within and across individuals, offers a systematic organization of associations.

All statistical analyses were performed in R. Linear mixed models were fit with the function lmer and generalized linear mixed models were fit with the function glmer, from the package lme4 (v1.1.38) (Bates et al., 2015). Estimated marginal means were computed with the package emmeans (v2.1.0) (Lenth and Piaskowski, 2026). To reduce auto-correlation, we used bouts instead of vocalizations as the basis for some analyses (see below for details).

### *10 Analysis 1: Per-hour analysis*

We analyzed the distribution of vocalizations and behaviors with respect to coarse contextual factors: hour of day, social category (breeding male, breeding female, helper), and group composition (number of adults, number of crow chicks, and the age of chicks).

#### Data

For each family on each day, we split each of the video and audio data into one-hour intervals and kept intervals starting between 4:00 and 20:00 UTC according to the sensor time. For each hourlong interval, we determined a list of individuals who had data covering at least 97% of the hour. Then, for each hour and individual present, we measured a suite of variables summarizing the vocalizations and behaviors of each individual. We measured the duration of time spent in the following states: any visit, any flight, low activity, and high activity. For each individual and vocalization type, we counted the number of vocalization bouts that started in the hour interval. When the appropriate sensors were not available, we recorded unknown. Overall, each observation consisted of the start hour of the measurement interval, the individual, their family, the individual's social category (breeding female, breeding male, or helper), the behavioral state or vocalization type measured, the corresponding duration or count in the hour, the number of crow chicks at the nest during this interval, the age of chicks in days during this interval, and the number of adults in the family. Overall, this procedure resulted in a dataset of 345,377 observations, including 57 individuals, with the number of observations per individual ranging from 90 to 14560. Individuals without audio-loggers only contributed observations of nest visits.

#### Model/statistics

For each vocalization type, we fit a Poisson mixed-effects regression model to predict the number of bouts in the hour. We included fixed effects for the hour of day, social category, number of adults, number of crow chicks, and the age of chicks. Hour of day was modeled categorically while number of adults, number of crow chicks, and the age of chicks were modeled linearly. We included random intercepts for each family and individual. For missing values, we added is_na dummy fixed effect (sex_is_na, role_is_na, n_crowchicks_is_na, chick_age_is_na). We only included observations that occurred in hours for which at least one count was observed. For each state, we fit a Gaussian linear mixed-effects regression model to predict the duration of time spent in the state, in that hour. We used the same fixed and random effects.

We obtained 95% confidence intervals for all fixed effects (hour of day: Figure S1, group composition: Figure S3). Estimated marginal means were also computed. For hour of day variables, we computed the proportionally-weighted deviation-from-mean ("eff") contrasts across hours. We calculated Benjamini-Hochberg adjusted $p$-values across these contrasts for vocalization types (number of comparisons = 629) and separately for behavioral states (number of comparisons = 118). In Figure S1 and Figure 5A, we plotted asterisks for hours with a significant positive deviation from the grand mean. For social category, we computed the equal-weighted means for each category and the pairwise contrasts between these means (Figure S2). We calculated Benjamini-Hochberg adjusted $p$-values across all pairwise social category contrasts (number of comparisons = 147). For group composition variables, we obtained $p$-values via Satterthwaite's degrees of freedom method using the function `as_lmerModLmerTest` in the lmerTest package v3.1.3 (Kuznetsova, Brockhoff, and Christensen, 2017). Then, we calculated Benjamini-Hochberg adjusted $p$-values across the fixed effects (Figure S3, number of comparisons = 147).

#### *11 Analyses 2 and 3: State-based analyses*

We investigated how vocalizations coincided with behavioral states (i.e., ongoing activities defined as intervals of time). This analysis identified scenarios typical of the production of vocalizations of different types, even if these are not tightly linked in time to discrete events like the beginning or end of a flight. First, we investigated how the type of a behavioral state predicts the rate of vocalization production. Second, we investigated how the type of a bout of vocalizations predicts the concurrent behavioral state.

##### Data

We considered the following mutually exclusive behavioral states: arriving flight, leaving flight, short flight, long flight, alternated visit, repeated visit, low activity and high activity. When the appropriate sensors were not available to determine a known vocalization count or behavioral state, we recorded unknown and did not include any observation.

First, for each behavioral state interval, we call the individual performing that behavior the *behaving individual*. We computed the rate of vocalizations made by the behaving individual (Figure S5, top-left) as well as by other individuals from the same family (Figure S5, bottom-left). Specifically, we recorded the duration of the behavioral state, and counted the number of calls of each vocalization type made by each tagged bird in the family. We also counted the number of calls made by crow chicks that were recorded on the behaving bird's audio-logger. Each observation consisted of the behaving individual, the vocalizing individual (which could be identical to the behaving individual), the type of the behavioral state, the type of the vocalization, the date and time of the start and end of the behavioral state, and the vocalization count. We filtered to behavioral intervals that did not include nighttime (starting between 04:00 and 20:00 UTC sensor time and ending between 04:00 and 21:00 UTC sensor time).

Overall, this procedure resulted in a dataset of 284,934 intervals in which the behaving and vocalizing individuals were the same, with 43 individuals represented. The least common state was repeated visit with 939 intervals and the most common state was high-activity with 134525 intervals. It resulted in a dataset of 196747 intervals in which the focal and vocalizing individuals were different, with 53 pairs of individuals represented. Again, the least common state was repeated visit with 928 intervals and the most common state was high-activity with 99102 intervals.

Second, for each vocalization bout, we recorded the behavioral state during which it began, for the vocalizing individual (Figure S5, top-right) as well as the other tagged birds from the same family (Figure S5, bottom-right). Each observation consisted of the vocalizing individual, the behaving individual (which could be identical to the vocalizing individual), the type of the vocalization bout, the type of the behavioral state, and the date and time of the start of the vocalization bout. We filtered to vocalization bouts which started during 04:00 and 20:00 UTC sensor time.

Overall, this procedure resulted in a dataset of 28535 bouts when the vocalizing and behaving individual were the same, with 32 individuals represented. The least common call type was E36 with 8 bouts, the most common call type was E4 with 3770 bouts, and the median number of bouts was 390. It also resulted in a dataset of 20652 bouts when the behaving and vocalizing bird were different, with 49 pairs of individuals represented. The least common call type was E47 with 2 bouts, the most common call type was E4 with 3843 bouts, and the median number of bouts was 302.

##### Model/statistics

First, we conditioned on the occurrence of a behavioral state to predict the rate of vocalizations of a certain type. For each vocalization type, we fit a Poisson mixed-effects regression model to predict the vocalization rate during the behavioral interval. We included a fixed effect for the state type, an offset corresponding to the log duration of the state, and random intercepts for the family, behaving individual, and date+hour of day. When a state type had no positive counts for a vocalization type, we omitted it from the model predicting counts for that type (this enabled the models to converge during training; omissions are indicated in Figure S5 by striped red blocks). We fit two models per call type, one for when the vocalizing bird is the behaving bird (Figure S5, top-left) and one for when they are different (Figure S5, bottom-left). In addition to a prediction for each vocalization type separately, we also fit a model to predict the count of all vocalizations (rows labeled ‘total’). In total, we fit 85 models (two for each vocalization type, plus two ‘total’ outcomes and one ‘crow chicks’ outcome).

Estimated marginal means were computed. To express rates per minute, we set the offset to the log of sixty seconds, and computed the equal-weighted deviation-from-mean (“eff”) contrasts for the state type levels. Figure S5 (left) reports the differences in log rates. Each row of the left-side of Figure S5 corresponds to one deviation-from-mean computation across the states. We adjusted the $p$-values with the Benjamini-Hochberg correction, with the number of comparisons equal to the number of states summed over the vocalization outcomes (# comparisons = 525), covering when the vocalizing and behaving birds are the same and when they are different.

For conditioning on the occurrence of a vocalization to predict the concurrent state, we fixed the state of interest and set the outcome to be a binary variable indicating whether the bird was in that state or not. For each state, we fit a linear mixed-effects regression model to predict the binary outcome variable. We included a fixed effect for the vocalization type (number of levels = 41) and random intercepts for the family, behaving individual and date+hour of day. We fit two models per behavioral state, one for when the vocalizing bird is the behaving bird (Figure S5, top-right) and one for when they are different (Figure S5, bottom-right). We attempted to use a logistic regression model, but this was difficult to fit due to the high proportion of zero entries. We note the precedence for using linear models rather than non-linear models when predicting binary variables (Gomila, 2021; Hellevik, 2009), and that one benefit of using a linear mixed model is that the effects are interpretable as probabilities. In total, we fit 16 models (two models for each state).

Estimated marginal means were computed. We computed the equal-weighted deviation-from-mean (“eff”) contrasts for the vocalization type levels. Each column of the right-side of Figure S5 (separately for top and bottom), corresponds to one deviation-from-mean computation across call types. We adjusted the $p$-values with the Benjamini-Hochberg correction, with the number of comparisons equal to the number of states times the number of call types times two (accounting for whether the vocalizing and behaving bird are the same).

### *12 Analysis 4: Event-based analysis*

We next analyzed vocalizations and behaviors as events. Events are operationalized as a discrete timepoint. In our dataset, they represented changes in behavioral states, such as the beginning or end of a flight. We looked at whether the occurrence of a *reference event* (equivalent to the *precedent event*) of a particular type was associated with the next *outcome event* of a particular type (equivalent to the *subsequent event*) occurring sooner or later than expected on average. In this analysis, between the reference and outcome events, there can be any number of intervening events (of a different type to the outcome event).

#### Data

As events, we considered the start and end times of the states which were considered in the state-based analysis (16 event types), and an additional 41 event types corresponding to the vocalization types, resulting in T=57 event types in each analysis. When relating vocalizations and behaviors, we used the beginning of vocalization bouts of each type as the vocalization events. When relating vocalizations to vocalizations, we used the start of single calls of each type as the vocalization events.

For each reference event of a type, we recorded the duration until the next outcome event of each type. These durations were recorded separately for events recorded by the reference individual, and for events recorded by other individuals in the family. For some pairs of events, there were periods of missing data between the occurrences of the events. For example, between a reference vocalization event and the outcome event corresponding to the start of the next known nest visit, there may be a period of time where no video data was recorded and therefore whether any nest visits occurred is unknown. To track this, we also recorded the summed duration of any missing data intervals between the reference and outcome events.

Therefore, each observation consisted of the reference individual, the outcome individual (which could be identical to the reference individual), the type of the reference event, the type of the outcome event, the date and time of the reference event, the time between the reference and outcome events (in seconds), and the duration of missing data between the reference and outcome events (in seconds). We excluded any observation with the duration of missing data greater than or equal to 60 seconds. We excluded observations if the reference event did not occur between 04:00 and 20:00 UTC.

When relating behaviors to vocalizations, this procedure resulted in a dataset of 584,606 reference events when the reference and outcome individuals were the same (54 individuals). For behavior precedents, the end of the visit repeated was the least common with 1430 events, the start of the high activity period was the most common with 129359 events, and the median was 6098. For vocalization precedents, E47 was the least common with 61 events, E4 was the most common with 8707 events, and the median was 825.5. It also resulted in a dataset of 1435741 reference events when the reference and outcome individuals were different (76 pairs). For behavior precedents, the end of repeated visits was the least common with 2787 events, the start of high activity periods was the most common with 327083 events, and the median was 15766. For vocalization precedents, E36 was the least common with 29 events, E4 was the most common with 16418 events, and the median was 1682.

When relating vocalizations to vocalizations, this procedure resulted in a dataset of 113964 reference events when the reference and outcome individuals were the same (43 individuals). E47 was the least common with 119 events, E4 was the most common with 18150 events, and the median was 1497. It also resulted in a dataset of 209195 reference events when the reference and outcome individuals were different (63 pairs). E36 was the least common with 32 events, E4 was the most common with 34467 events, and the median was 2710.5.

#### Model/statistics

For each type of outcome event $T$, we fit a Gaussian mixed-effects regression model to predict the log-transformed time between a reference event and the outcome event of type $T$. We included a fixed effect for the type of the reference event, and random intercept for the individual, family, date+hour of day. We used one model for the vocalizations as reference events (number of levels = 41; Figures S6/S7, left) and a different model for the other behavioral events (number of levels = 16; Figures S6/S7, right). In total, we fit 228 models (four for each type of outcome event: matching vs. different birds times vocalization vs. behavioral reference events).

Estimated marginal means were computed. We computed the equal-weighted deviation-from-mean ("eff") contrasts for the precedent event type levels. Each row in each subpanel of Figures S6-S7 corresponds to a deviation-from-mean computation across precedent events. We adjusted the $p$-values with the Benjamini-Hochberg correction separately for models with behavior reference events (# comparisons = 1824) and models with vocalization reference events (# comparisons = 4674).

We note that the relationships between a crow's behavioral events (Figure S6; behavior $\rightarrow$ self behavior, top left) provided a built-in validation for the analysis. The start of behavioral states anticipated the end of the same state, relative to other events. We also see replications of the state-based analyses. For example, E4 is strongly associated with the signaller being at the nest (Figure S5): here, it is preceded by nest arrivals and inhibited after nest departures (Figure S6; behavior $\rightarrow$ self vocalization, bottom-left).

#### Robustness check

We defined a binary outcome variable of whether an outcome event happened within the ten minutes after the reference event. For each type of outcome event $T$, we fit a linear mixed-effects regression model to predict the binary outcome variable (Gomila, 2021; Hellevik, 2009). As in our original model specification, we included a fixed effect for the type of the reference event, and random intercept for the individual, family, date+hour of day. We found that the results were largely consistent between the two model specifications (Figure S12).

### Supplementary Figures

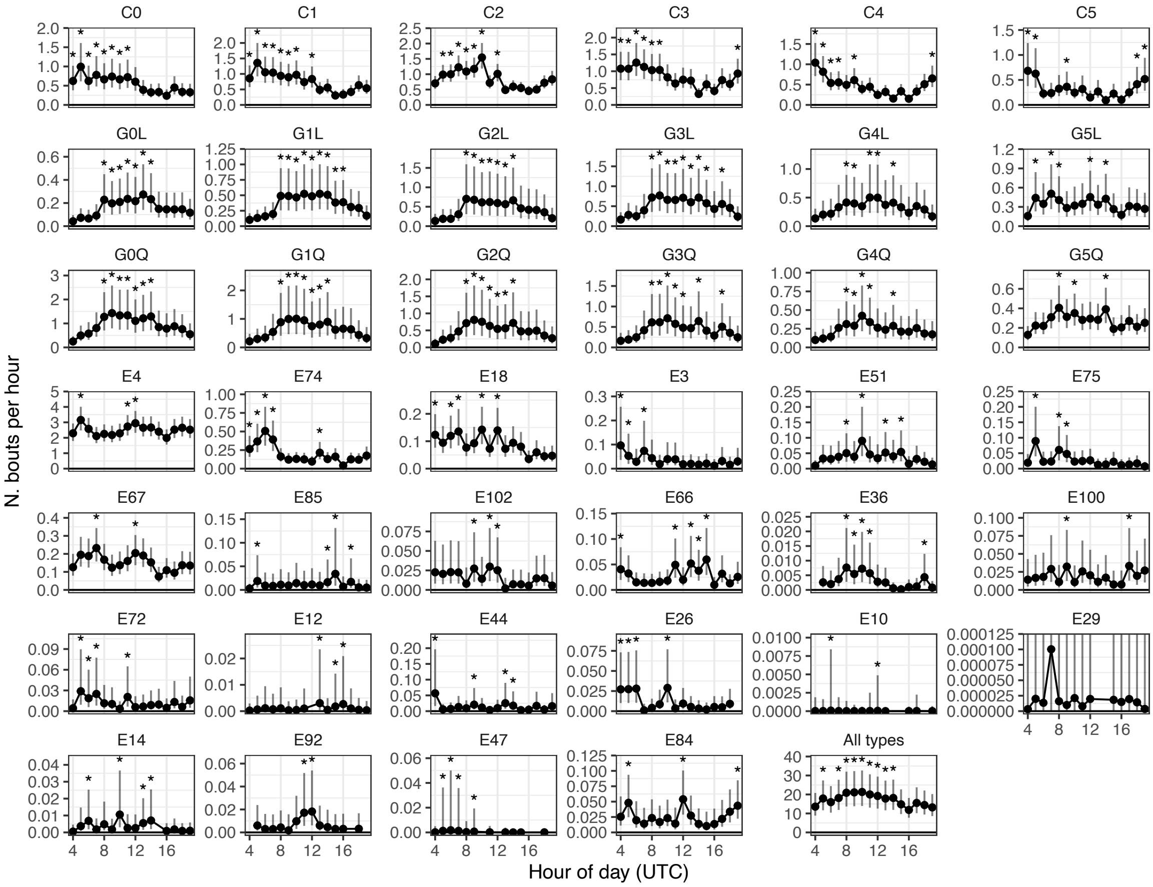


Figure S1. **Time-of-day results from analysis 1.** Call types show varied profiles over the day. Model-based means for the number of bouts produced by an individual in an hour, for each call type. Error bars show 95% CIs, asterisks show hours that had higher rates than the model-based mean at $p_{\mathrm{BH}} < 0.05$. Exceptional calls are ordered by their proportion in the dataset.

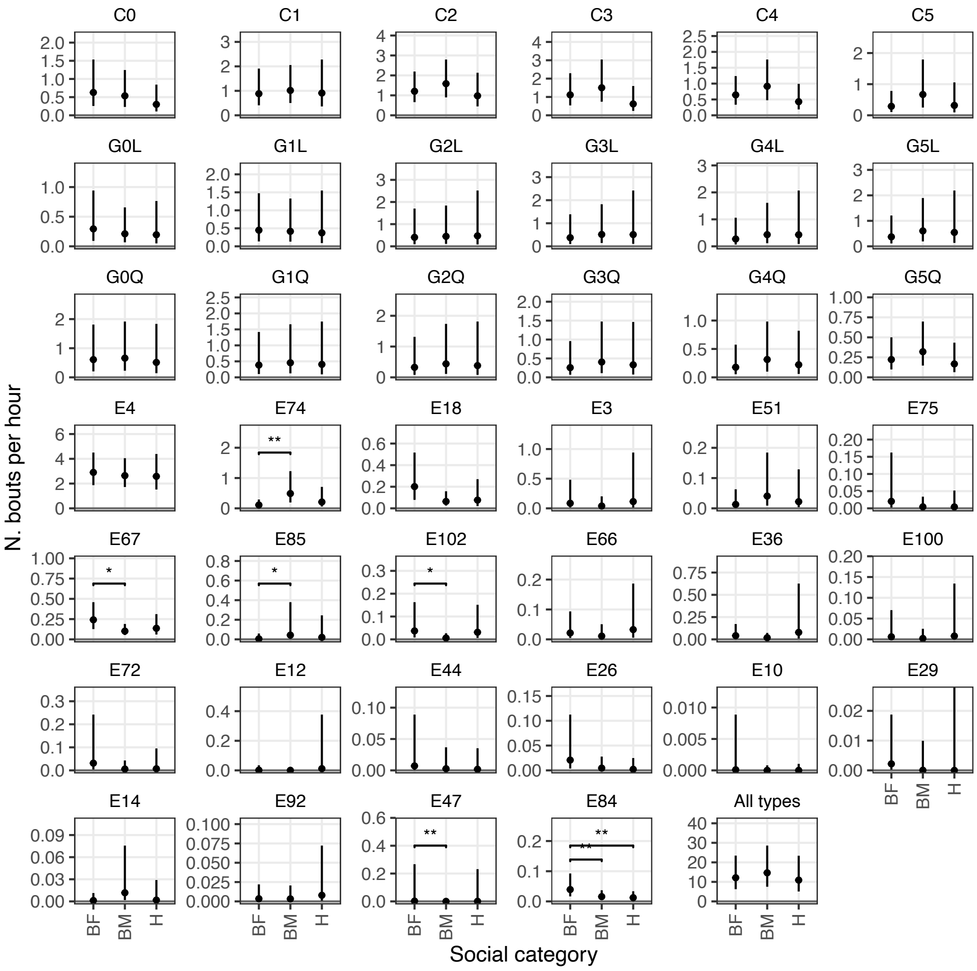

Figure S2. **Social-category results from analysis 1.** Many call types are shared by crows of different social categories. For each call type, plot shows model-based means for the number of bouts per hour by social category. Social categories: BF (Breeding female), BM (Breeding male), H (Helper). Asterisks shows significance in pairwise comparisons: * $p_{\text{BH}} < 0.05$, ** $p_{\text{BH}} < 0.01$. Exceptional calls are ordered by their proportion in the dataset.

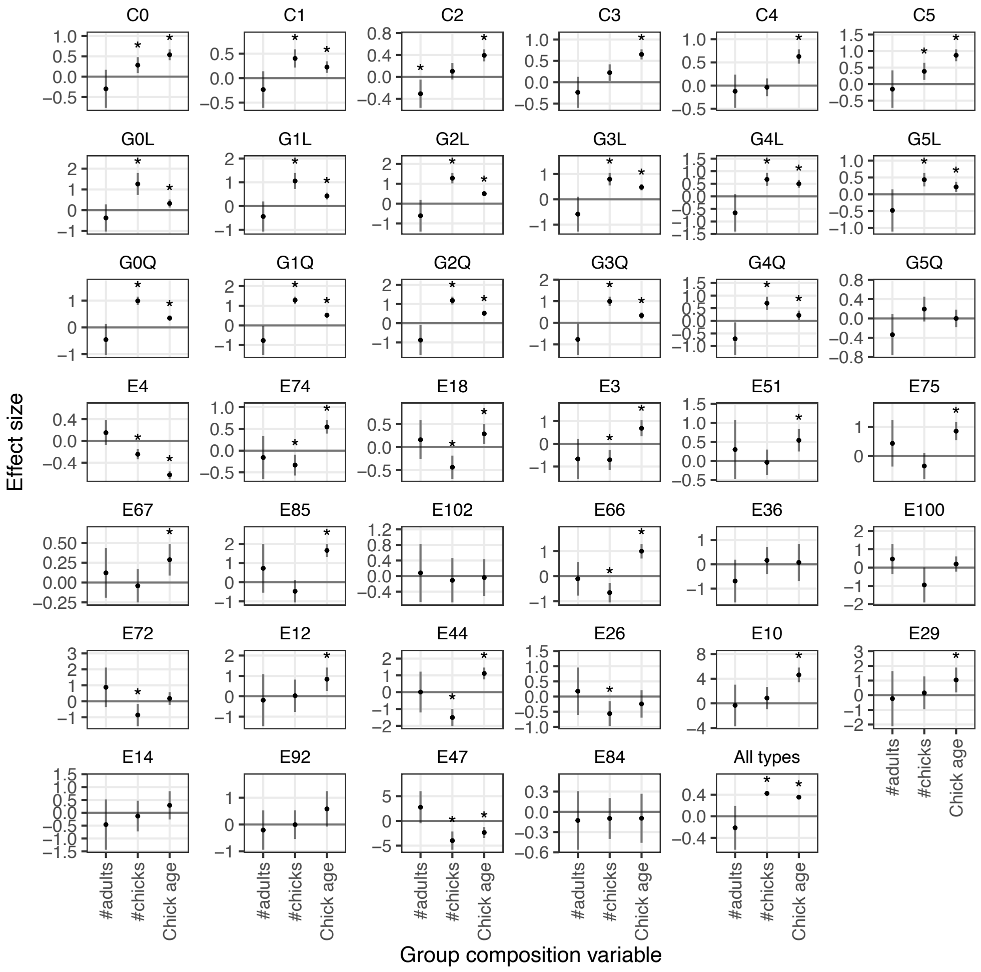


**Figure S3**. **Group composition results from analysis 1.** The number of crow chicks and their age is a common modulator for the usage of several vocalization types. Plots show the effect size of group composition variables on the number of bouts per hour. Asterisks show $p_{\mathrm{BH}} < 0.05$. Exceptional calls are ordered by their proportion in the dataset.

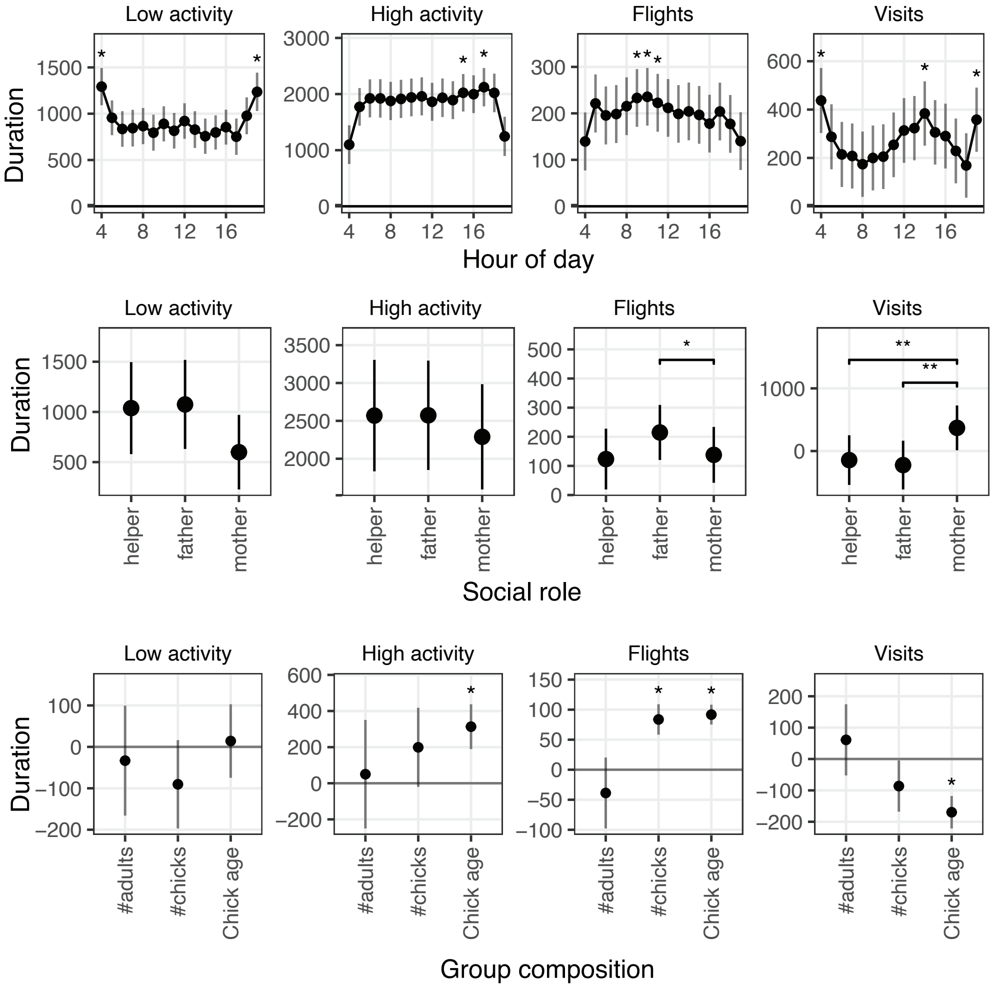


Figure S4. **Analysis 1, where outcome variables are the duration of behavioral states in an hour.** Plotting follows Figures S1-S3.

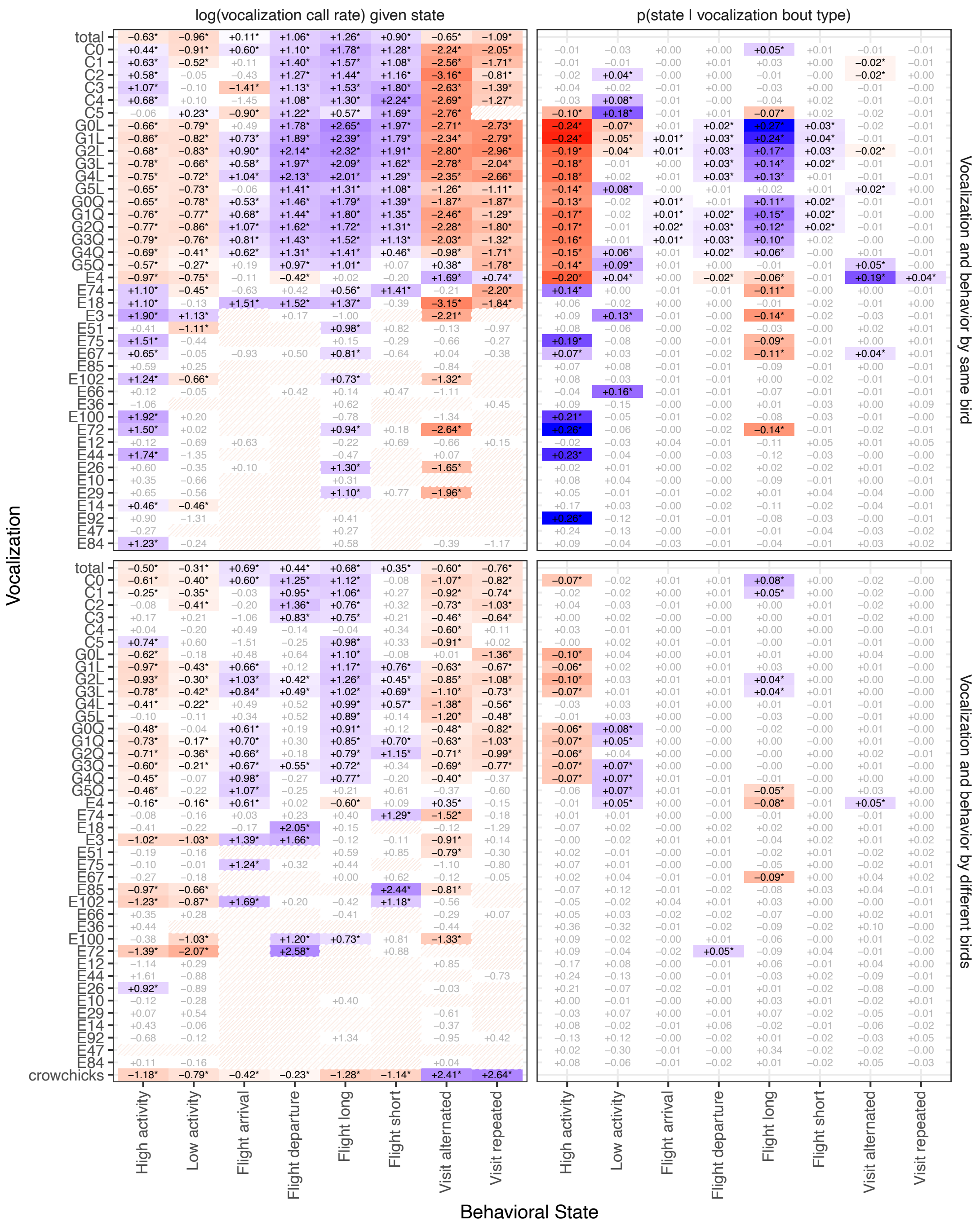


**Figure S5. Association matrix for Analyses 2-3 (state-based).** Left: States concurrent with vocalization production. Vocalization rate varied based on the concurrent behavioral state of the signaller, or of potential receivers in the same group. Top: vocalizations produced by the signaller, given the signaller's behavioral state. Bottom: vocalizations produced by the signaller, given the behavioral state of another individual. Each row shows the deviation from the equal-weighted, model-based mean of the log rate (vocalizations/minute) in each state. Blue/red entries = higher/lower than the model-based mean respectively, with $p_{\mathrm{BH}} < 0.05$; greyed entries indicate non-significant contrast. Striped entries indicate that this state had no positive counts of the target vocalization, and was not included in the model. Exceptional calls are sorted based on their proportion in our dataset. Right: Concurrent states inferred from vocalizations. Vocalization bout type affects the probability of whether a signaller or potential receivers were concurrently in a given behavioral state, or not. Top: what could be inferred about what a signaller was doing, given their vocalization. Bottom: what could be inferred about the state of receivers, given a signaller's vocalization. Each column shows the deviation from the equal-weighted, model-based mean of the probability of being in the state vs. not being in that state, for each vocalization type. Color scheme as in left panels.

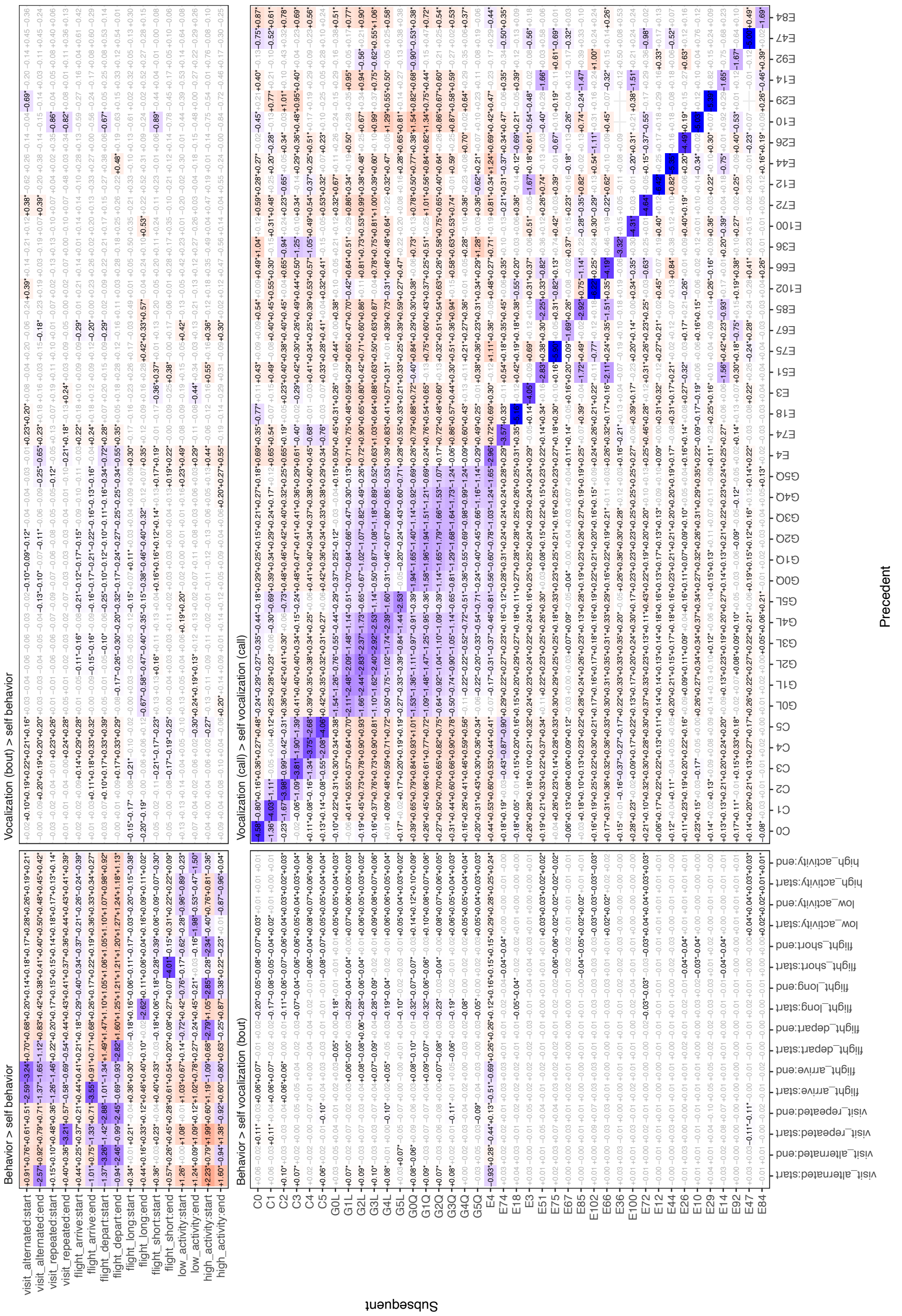


Figure S6. **Association matrix for Analysis 4 (event-based), where events are performed by the same bird (rotate to view).** Within a quadrant, each row shows the deviation from the equal-weighted, model-based mean of the log time (seconds) until the subsequent event, for each type of precedent event. Blue/red entries = sooner/later than the model-based mean respectively, significant after multiple comparisons correction. Top-left quadrant: Behavioral events preceding behavioral events. Top-left quadrant: Vocalization bouts preceding behavioral events. Bottom-left quadrant: Behavioral events preceding vocalization bouts. Bottom-right quadrant: Vocalizations preceding vocalizations (using single calls, rather than bouts). Exceptional calls are sorted based on their proportion in our dataset.

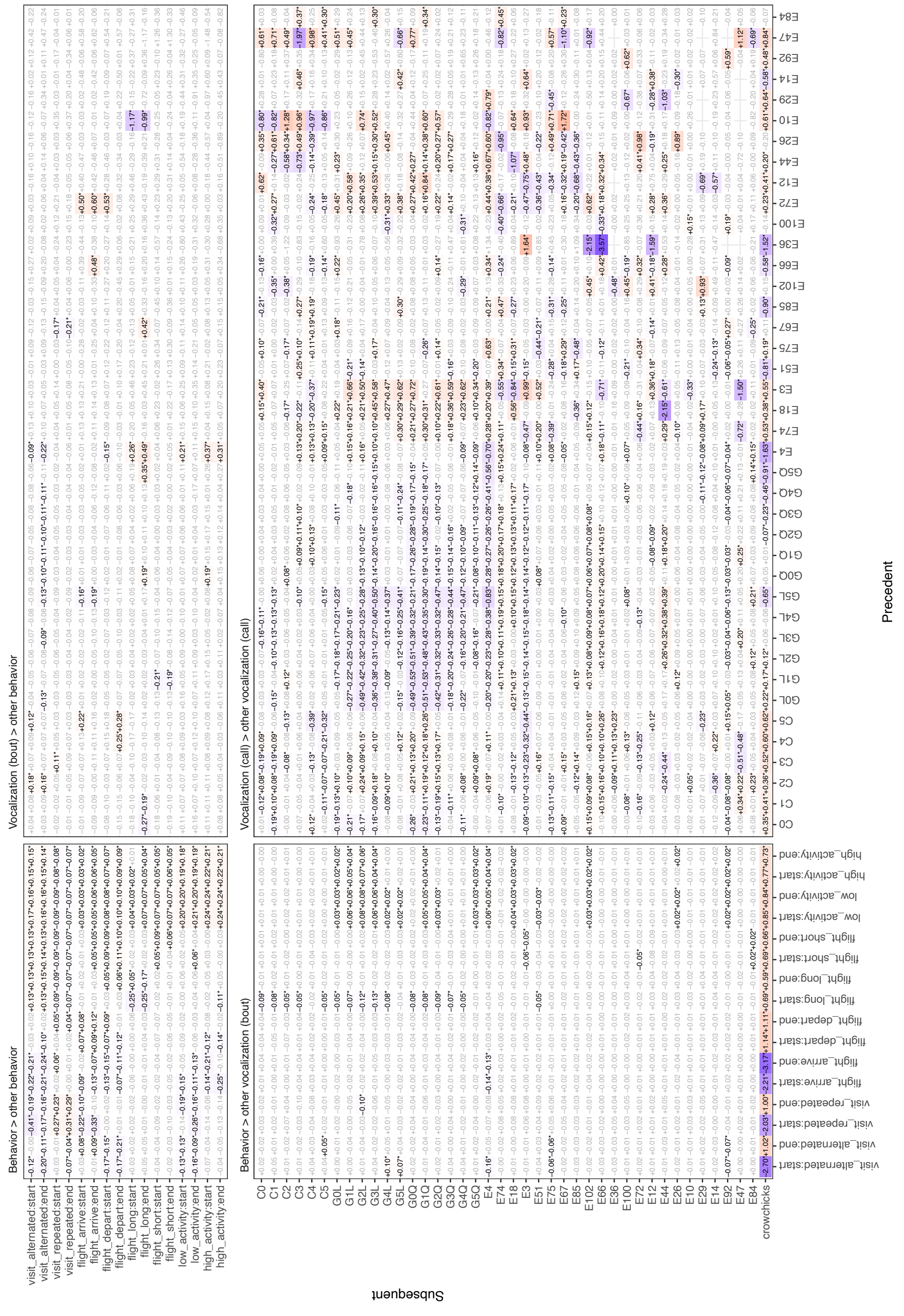


**Figure S7. Association matrix for Analysis 4 (event-based), where events are performed by different birds (rotate to view).** Same color scheme as Figure S6.

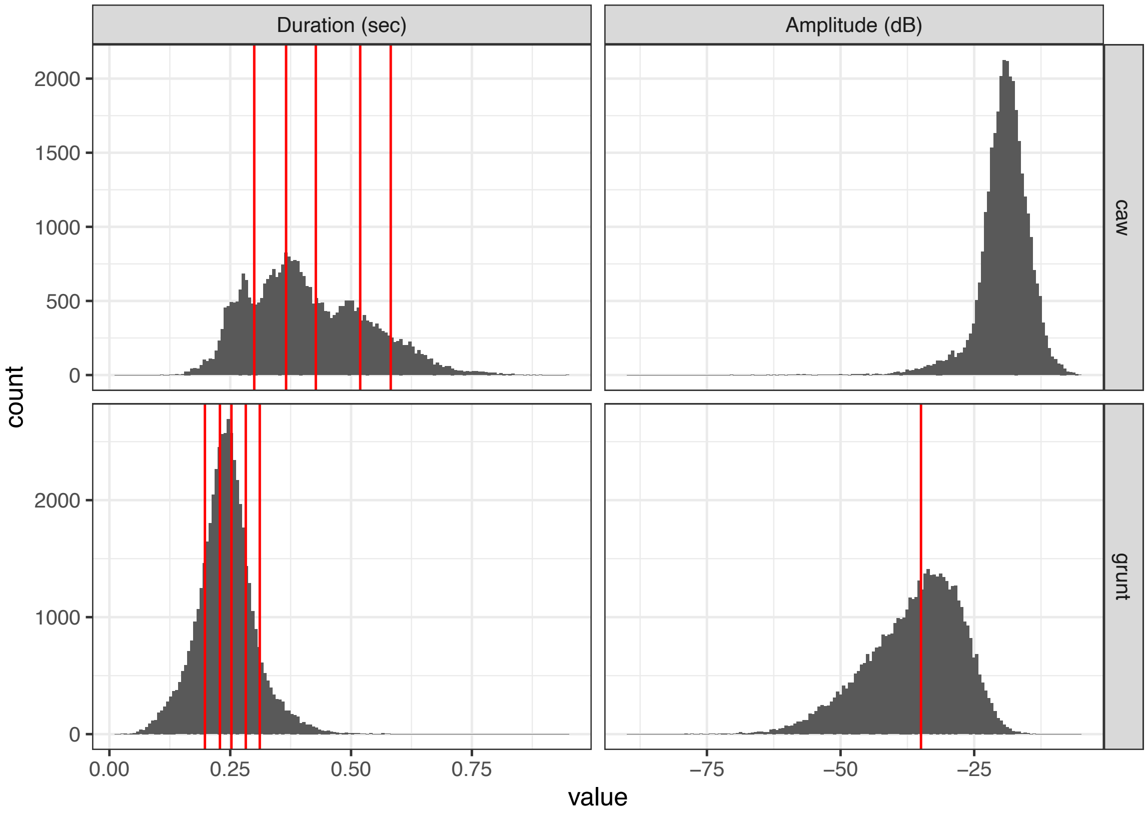


Figure S8. **Acoustic features of grunts and caw super-categories.** Quantile boundaries used to split the super-categories are shown in red.

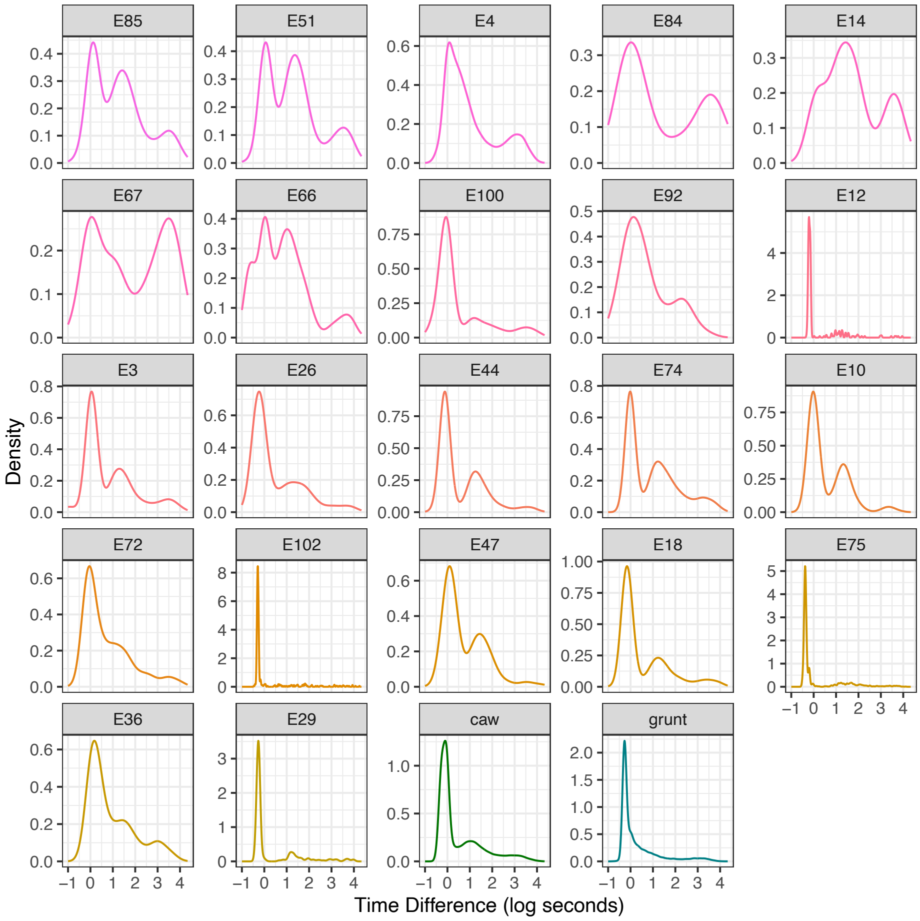


Figure S9. **Kernel density estimates for the timing between calls of the same type, by call type**. Caws and grunts are kept as super-categories. We used the duration of first valley, taking the median across call types, to define the bout threshold. Exceptional call types are ordered by mean amplitude.

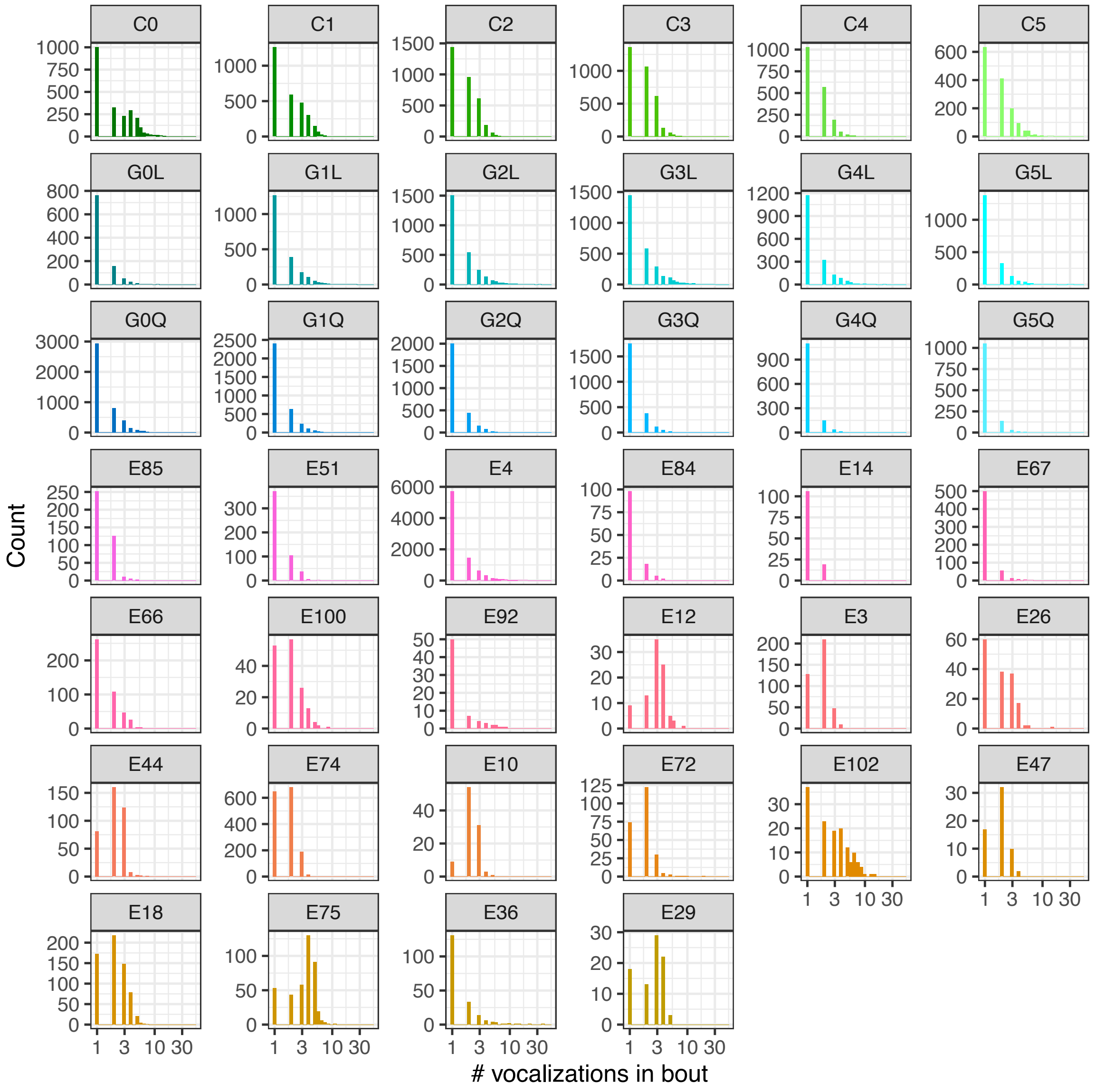


**Figure S10. Distribution of number of vocalization in a bout, per call type.** Colors follow Figure 2, with exceptional calls ordered in terms of mean amplitude.

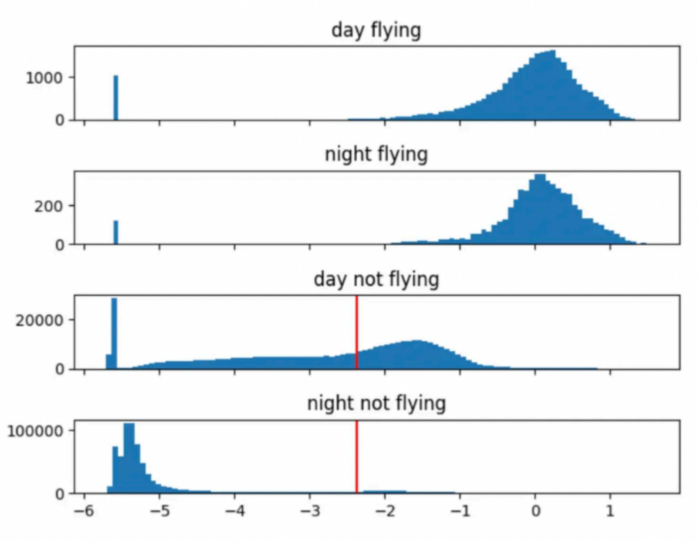


Figure S11. **Distribution of overall dynamic body acceleration.** Distribution of overall dynamic body acceleration, averaged in one-second bins. X-axis shows units of $g$ on a log scale; y-axis shows bin counts. Red line shows the median value for not flying bins during the day. The magnitude during flight is highest. The magnitude of not-flying at night is the lowest, interpreted as the birds staying still while they are asleep. The magnitude of not-flying during the day is in the middle, with a long left skew.

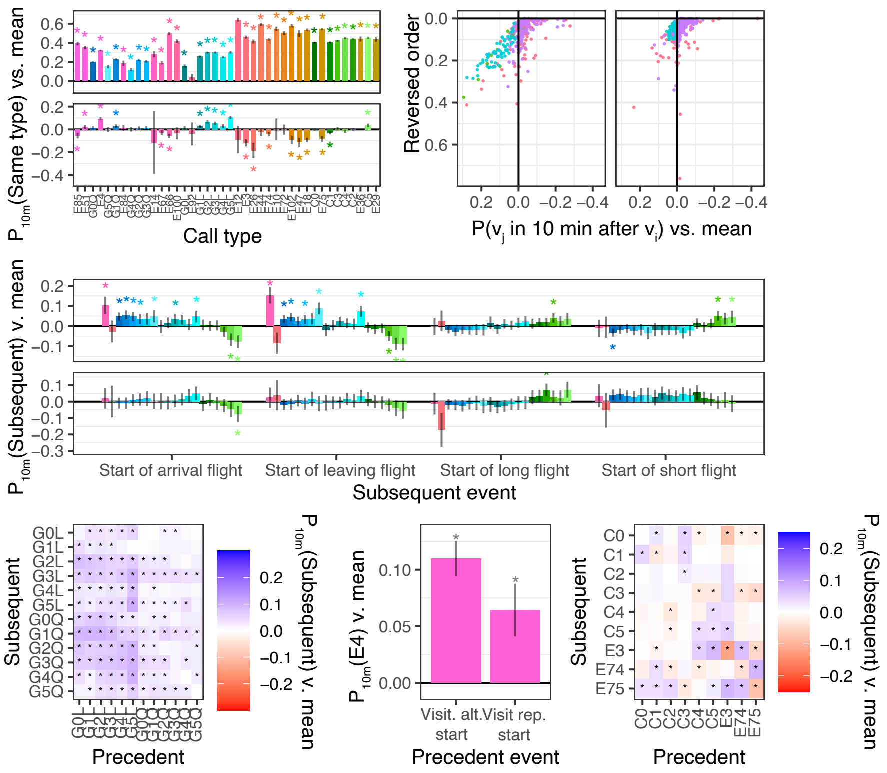


Figure S12. **Analysis 4 results are largely consistent with an alternative model specification.** Figures 2D, 2E, 3E, 3D, 4B, 5D replotted using alternative binary specification of event-based analysis (see Materials and Methods, Section 12). $P_{10m}$(subsequent event) indicates the probability that the subsequent event occurs in the ten minutes after the precedent event.

| Year | Territory | Number of adults | Number of adults with biologgers | Duration of synchro-nized video (hours) | Number of crow chicks at beginning of video annotation period | Chick deaths during video annotation period | Number of cuckoo chicks |
|---|---|---|---|---|---|---|---|
| 2018 | AW | 4 | 2 | 19.1 | 1 | 0 | 0 |
| 2018 | AY | 3 | 2 | 26.0 | 2 | 0 | 0 |
| 2018 | BC | 3 | 3 | 12.0 | 1 | 1 | 0 |
| 2018 | CT | 3 | 1 | 13.8 | 3 | 2 | 0 |
| 2018 | N1 | 2 | 1 | 16.0 | 2 | 0 | 0 |
| 2018 | O | 3 | 2 | 44.0 | 3 | 1 | 2 |
| 2018 | VAE | 2 | 2 | 3.0 | 2 | 0 | 0 |
| 2019 | AL | 4 | 3 | 113.0 | 1 | 0 | 1 |
| 2019 | BA | 4 | 2 | 89.0 | 4 | 0 | 0 |
| 2019 | BD | 3 | 2 | 32.0 | 2 | 0 | 0 |
| 2019 | BPBO | 2 | 1 | 19.0 | 2 | 0 | 0 |
| 2019 | BUCK | 5 | 3 | 64.0 | 3 | 0 | 0 |
| 2019 | BV | 5 | 2 | 93.0 | 5 | 0 | 0 |
| 2019 | CA | 3 | 1 | 5.0 | 2 | 0 | 0 |
| 2019 | CV | 2 | 2 | 36.8 | 3 | 0 | 0 |
| 2019 | CW | 2 | 1 | 70.0 | 1 | 1 | 0 |
| 2019 | EB | 3 | 2 | 37.0 | 3 | 0 | 0 |
| 2019 | N7 | 2 | 2 | 47.0 | 2 | 0 | 1 |
| 2021 | BPBO | 2 | 2 | 64.0 | 4 | 0 | 0 |
| 2021 | BQ | 2 | 1 | 49.0 | 1 | 1 | 0 |
| 2021 | EB | 3 | 1 | 0.0 | NA | NA | NA |
| 2021 | FA | 4 | 2 | 71.0 | 4 | 0 | 0 |
| 2021 | N4 | 2 | 2 | 70.0 | 2 | 0 | 0 |
| 2021 | LL | 5 | 2 | 0.0 | NA | NA | NA |

Table S1. **Summary of 24 social groups in dataset.**

Table S2. **Summary of individuals in dataset.** Crows without wing tags are not included here but supplied additional nest visits (labeled as "untagged crow").

| Year | Territory | Tag ID | Role | Sex | N. focal vocalizations | N. visits | N. flights | Dur. audio (h) | Approx. dur. functioning daytime audio (h) |
|---|---|---|---|---|---|---|---|---|---|
| 2018 | AW | B5 | H | M | 634 | 30 | 44 | 3.22 | 3.22 |
| 2018 | AW | A5 | B | F | 6978 | 30 | 1236 | 157.34 | 108.03 |
| 2018 | AW | B4 | NA | NA | 0 | 15 | 0 | 0.00 | 0.00 |
| 2018 | AY | A4 | H | F | 1777 | 38 | 481 | 79.42 | 59.43 |
| 2018 | AY | B1 | H | M | 190 | 53 | 50 | 6.43 | 0.08 |
| 2018 | BC | A0 | B | F | 6168 | 13 | 594 | 141.09 | 93.99 |
| 2018 | BC | C1 | H | M | 6601 | 19 | 1367 | 147.54 | 105.98 |
| 2018 | BC | B0 | B | M | 5275 | 11 | 854 | 145.55 | 103.18 |
| 2018 | CT | D6 | B | F | 3171 | 26 | 467 | 100.29 | 69.17 |
| 2018 | CT | A1 | B | M | 0 | 3 | 0 | 0.00 | 0.00 |
| 2018 | N1 | B6 | B | F | 393 | 12 | 76 | 26.54 | 20.01 |
| 2018 | N1 | A2 | B | M | 0 | 38 | 0 | 0.00 | 0.00 |
| 2018 | O | A7 | B | M | 221 | 83 | 118 | 7.36 | 5.55 |
| 2018 | O | B2 | B | M | 2872 | 111 | 1352 | 132.39 | 97.51 |
| 2018 | VAE | A6 | B | F | 990 | 5 | 348 | 56.53 | 40.19 |
| 2018 | VAE | A3 | B | M | 1968 | 10 | 688 | 58.90 | 38.74 |
| 2019 | AL | K8 | B | M | 4598 | 145 | 1639 | 154.96 | 116.38 |
| 2019 | AL | M9 | H | F | 3847 | 101 | 1018 | 160.44 | 117.88 |
| 2019 | AL | L8 | B | F | 1492 | 170 | 536 | 164.42 | 60.11 |
| 2019 | BA | E3 | B | F | 2024 | 279 | 1709 | 139.74 | 101.48 |
| 2019 | BA | F4 | B | M | 0 | 175 | 0 | 0.01 | 0.01 |
| 2019 | BD | C5 | B | M | 572 | 59 | 132 | 5.67 | 4.72 |
| 2019 | BD | E1 | B | F | 808 | 61 | 412 | 45.78 | 30.75 |
| 2019 | BPBO | L1 | H | F | 2262 | 30 | 583 | 49.83 | 34.99 |
| 2019 | BUCK | M5 | B | F | 4164 | 109 | 1388 | 152.68 | 79.00 |
| 2019 | BUCK | N8 | B | M | 758 | 45 | 817 | 108.05 | 53.68 |
| 2019 | BUCK | L5 | H | F | 678 | 115 | 517 | 116.57 | 79.89 |
| 2019 | BUCK | K6 | H | F | 0 | 32 | 0 | 0.00 | 0.00 |
| 2019 | BV | N5 | H | M | 1662 | 266 | 813 | 109.67 | 69.54 |
| 2019 | BV | L4 | B | M | 1500 | 355 | 1914 | 146.37 | 104.05 |
| 2019 | BV | B3 | NA | NA | 0 | 8 | 0 | 0.00 | 0.00 |
| 2019 | CA | M8 | B | F | 229 | 17 | 96 | 194.87 | 5.87 |
| 2019 | CV | B8 | B | M | 1772 | 76 | 653 | 50.46 | 37.23 |
| 2019 | CV | A8 | B | F | 1593 | 59 | 297 | 52.41 | 37.19 |
| 2019 | CW | E4 | B | F | 4911 | 233 | 1289 | 137.71 | 98.38 |
| 2019 | EB | B4 | H | M | 649 | 68 | 307 | 53.42 | 37.80 |
| 2019 | EB | A9 | B | F | 1246 | 37 | 299 | 46.62 | 31.10 |
| 2019 | N7 | P3 | B | F | 2312 | 52 | 908 | 86.57 | 61.99 |
| 2019 | N7 | E5 | B | M | 1703 | 131 | 885 | 98.76 | 74.18 |
| 2021 | BPBO | L1 | B | F | 4578 | 163 | 2008 | 124.51 | 89.46 |
| 2021 | BPBO | B9 | B | M | 5517 | 284 | 1061 | 82.37 | 63.59 |
| 2021 | BQ | J7 | B | F | 5929 | 94 | 1070 | 141.18 | 100.70 |

| | | | | | | | | | |
|---|---|---|---|---|---|---|---|---|---|
| 2021 | BQ | B9 | B | M | 0 | 43 | 0 | 0.00 | 0.00 |
| 2021 | EB | E9 | B | M | 2031 | 0 | 491 | 65.99 | 46.94 |
| 2021 | FA | F6 | B | F | 5112 | 230 | 1594 | 136.20 | 97.45 |
| 2021 | FA | E7 | B | M | 1270 | 177 | 782 | 95.81 | 67.33 |
| 2021 | FA | H6 | NA | NA | 0 | 119 | 0 | 0.00 | 0.00 |
| 2021 | N4 | J6 | B | M | 6218 | 202 | 1498 | 124.97 | 88.12 |
| 2021 | N4 | H9 | B | F | 162 | 109 | 22 | 156.14 | 0.01 |
| 2021 | LL | H5 | B | F | 3652 | 0 | 1453 | 132.02 | 92.39 |
| 2021 | LL | E6 | B | M | 4189 | 0 | 2234 | 153.50 | 109.31 |

| Call type | N. calls | N. unique individual | Proportion by most represented individual | Shannon diversity index | N. effective individuals (Shannon) | N. effective individuals (Simpson) |
|---|---|---|---|---|---|---|
| grunt | 49427 | 43 | 0.093 | 3.348 | 28.448 | 22.859 |
| G0L | 1458 | 39 | 0.175 | 3.029 | 20.677 | 13.893 |
| G1L | 4110 | 40 | 0.165 | 3.070 | 21.532 | 15.156 |
| G2L | 5812 | 41 | 0.150 | 3.128 | 22.825 | 16.729 |
| G3L | 6586 | 41 | 0.160 | 3.156 | 23.483 | 16.908 |
| G4L | 3340 | 39 | 0.105 | 3.176 | 23.944 | 18.245 |
| G5L | 3407 | 41 | 0.140 | 2.983 | 19.740 | 13.572 |
| G0Q | 8428 | 43 | 0.198 | 3.125 | 22.756 | 14.542 |
| G1Q | 5775 | 42 | 0.133 | 3.268 | 26.248 | 20.048 |
| G2Q | 4073 | 41 | 0.100 | 3.295 | 26.980 | 21.682 |
| G3Q | 3299 | 41 | 0.096 | 3.278 | 26.535 | 20.842 |
| G4Q | 1603 | 40 | 0.140 | 3.104 | 22.296 | 15.331 |
| G5Q | 1536 | 43 | 0.144 | 3.153 | 23.407 | 16.222 |
| caw | 33427 | 43 | 0.100 | 3.351 | 28.533 | 22.951 |
| C0 | 6686 | 40 | 0.177 | 3.009 | 20.258 | 13.740 |
| C1 | 6685 | 41 | 0.115 | 3.200 | 24.529 | 18.903 |
| C2 | 6685 | 41 | 0.087 | 3.295 | 26.967 | 22.543 |
| C3 | 6685 | 43 | 0.136 | 3.218 | 24.969 | 18.340 |
| C4 | 3343 | 39 | 0.095 | 3.263 | 26.132 | 20.321 |
| C5 | 3343 | 39 | 0.164 | 3.072 | 21.581 | 15.269 |
| E4 | 18369 | 43 | 0.103 | 3.353 | 28.597 | 21.753 |
| E74 | 2669 | 38 | 0.372 | 2.425 | 11.301 | 5.816 |
| E18 | 1510 | 35 | 0.236 | 2.520 | 12.424 | 8.495 |
| E75 | 1488 | 26 | 0.526 | 1.855 | 6.390 | 3.311 |
| E44 | 840 | 27 | 0.662 | 1.264 | 3.540 | 2.128 |
| E66 | 756 | 33 | 0.427 | 1.776 | 5.903 | 3.300 |
| E3 | 740 | 28 | 0.153 | 2.728 | 15.304 | 11.883 |
| E67 | 735 | 37 | 0.110 | 3.084 | 21.839 | 16.262 |
| E51 | 719 | 32 | 0.260 | 2.140 | 8.498 | 5.434 |
| E85 | 572 | 23 | 0.427 | 1.859 | 6.414 | 3.802 |
| E102 | 507 | 25 | 0.270 | 2.128 | 8.396 | 5.606 |
| E72 | 496 | 22 | 0.401 | 2.022 | 7.554 | 4.747 |
| E36 | 425 | 12 | 0.953 | 0.292 | 1.340 | 1.101 |
| E26 | 351 | 16 | 0.276 | 1.974 | 7.197 | 5.703 |
| E100 | 338 | 25 | 0.275 | 2.430 | 11.357 | 7.537 |
| E12 | 292 | 10 | 0.404 | 1.418 | 4.131 | 3.300 |
| E29 | 234 | 8 | 0.453 | 1.403 | 4.067 | 3.377 |
| E10 | 227 | 10 | 0.714 | 0.984 | 2.676 | 1.855 |
| E84 | 157 | 34 | 0.287 | 2.766 | 15.900 | 8.591 |
| E14 | 144 | 16 | 0.299 | 1.752 | 5.767 | 4.227 |
| E92 | 134 | 18 | 0.433 | 1.802 | 6.060 | 3.732 |
| E47 | 119 | 16 | 0.613 | 1.433 | 4.190 | 2.421 |
| total | 114676 | 43 | 0.061 | 3.459 | 31.790 | 27.120 |

Table S3. **Individuality metrics for each call type.**

**Table S4**. **Text description of call types**. Matches with prior works are uncertain because they are based on spectrograms and text descriptions alone.

| Our call type name | Description | Possible correspondence in Siriwardena | Possible correspondence in Cramp and Perrins |
|---|---|---|---|
| Caw | Typical crow "caw" sound. Stack of spectral bands, with maximum spectral energy between 1000 and 2000 Hz. Often amplitude modulated. Bands typically slope upwards at the beginning of the vocalization, and downwards at the end. Duration ranges from short (<0.2 sec, Caw0) to long (>0.7sec, Caw5). | Loud amplitude-modulated call | Advertising call |
| Grunt | A subdued "caw" sound. Compared to the Caw call type, lower amplitude, less prominent spectral bands, less pronounced amplitude modulation, less of a up-and-down frequency trajectory. Duration ranges from short (<0.1 sec, Grunt0Q/L) to medium (0.4 sec, Grunt1Q/L). In addition to amplitude differences, Q grunts have less pronounced spectral bands than L grunts | Low grunt calls | n/a |
| E3 | A brief, high-pitched accent note followed by a longer (>0.5 second) whine or honk. Both parts of the call have pronounced harmonics, with fundamental frequency approximately 1000Hz for the initial note and 500Hz for the remainder. | n/a | 'Motor-horn' variant of advertising call |
| E4 | A quiet growl, more subdued than a grunt. Noisy, with some variation on whether harmonic structure is apparent. Typically 0.3-0.5 sec. | n/a | n/a |
| E10 | A loud, descending, amplitude-modulated call (approx 0.5 sec), which is followed by a brief (<0.1 sec) accent note with the top of its spectral range extending higher. Both parts of the call have clear harmonic structure. | n/a | n/a |
| E12 | A honking call (approx 0.4 sec), with amplitude modulation and overall flat frequency contour. Most energy is concentrated in a single frequency band, around 1200Hz, although harmonics appear down to 200 Hz. | Loud amplitude-modulated call variant(?) | "keerk" variant of advertising call |
| E14 | A typically long (>0.5 sec) eerie "oo", with most energy in 500-700 Hz range. Spectral content can range from having one dominant component (a whistle), to sounding biphonated. Often preceded by a brief, high-pitched accent note and accompanied with bill-clacks. | n/a | n/a |
| E18 | A short (<0.1 sec) high-pitched harmonic stack followed by a longer (>0.3 sec.) descending, amplitude modulated tone. The first note has harmonic bands beginning around 400 Hz, and the second descends from there. | n/a | n/a |
| E26 | A rapid succession of two tones (<0.3 sec total), the first higher in pitch than the second. The first often longer than the second. | n/a | n/a |
| E29 | A high-pitched flat harmonic tone, approximately 0.3 sec long with peak energy around 1200Hz | n/a | n/a |
| E36 | A whiny, nasally call, 0.5 sec or longer. Often descending in frequency contour with pronounced harmonics. Little-to-no amplitude modulation. | n/a | n/a |
| E44 | A succession of two tones, up to 0.5 sec, with the opposite order of E26. The first note is lower-pitched than the second, while the second, shorter note is higher-pitched and tends to have more amplitude modulation. | n/a | n/a |
| E47 | A succession of two notes, around 0.7 sec. The first note is higher-pitched without amplitude modulation. The second, which is sometimes but not always longer than the first, is lower-pitched with amplitude modulation. Somewhat like E3 but typically with less dramatic jump in frequency contour and amplitude modulation on the second note. | n/a | n/a |
| E51 | A brief (<0.1 sec) low-frequency harmonic "urk" sound, often accompanied with bill clacks. Sometimes followed by a low-frequency pure tone of the same fundamental. | "Urk" call | n/a |
| E66 | A very brief, low-amplitude grunt (<0.1 sec) accompanied with bill clacks. Sometimes, but not always, it is followed by a longer, higher-pitched, amplitude-modulated trill (0.5 sec or more). The first note has harmonic bands going as low as 200 Hz. The second note, if it occurs, has most energy in the 600-800 Hz range. It may be dominated by a single component (a whistle) or have multiple. | n/a | n/a |

Table S4. **Text description of call types**. Matches with prior works are uncertain because they are based on spectrograms and text descriptions alone. *(continued)*

| Our call type name | Description | Possible correspondence in Siriwardena | Possible correspondence in Cramp and Perrins |
|---|---|---|---|
| E67 | Extremely brief (<0.05 sec) quiet “tut” or “zip” sound. | n/a | n/a |
| E72 | High-pitched trill, lasting 0.5-1 sec. Strong harmonic bands, with lowest around 1000Hz. Amplitude and frequency modulated, around an overall flat frequency contour. | n/a | n/a |
| E74 | Long (>0.7 sec), harsh call. No clear harmonic bands, with peak spectral energy around 1500-1700Hz. Flat pitch contour | Loud amplitude-modulated call variant(?) | Anxiety-call |
| E75 | Short (<0.2 sec), high pitched succession of two notes. The first has no amplitude modulation and strong harmonic bands beginning around 1100Hz. The second note has amplitude modulation and more apparent upper harmonics. Each note has a flat pitch contour. | n/a | Hawk-alarm call(?) |
| E84 | Short (<0.3) guttural “pa-choo” rattles, often descending in pitch. | n/a | n/a |
| E85 | Very short (0.1 sec), quiet grunts. Most spectral energy is around 400 Hz. Sometimes accompanied by bill clicks. | n/a | n/a |
| E92 | Percussive rattle calls, not guttural like E84. Ranges in length from 0.1-0.5 sec or longer. | Rattle | Rattle-call |
| E100 | High frequency, short (0.1 sec) ascending “zip” note. No clear harmonic bands, with most energy occurring in 600-1500 Hz range. | n/a | n/a |
| E102 | Like E18, but much shorter (<0.2 sec) and higher pitched, with harmonic bands starting around 1100 Hz for the first tone. | n/a | n/a |
